# The MYO1B and MYO5B motor proteins and the SNX27 sorting nexin regulate membrane mucin MUC17 trafficking in enterocytes

**DOI:** 10.1101/2023.03.06.530313

**Authors:** Sofia Jäverfelt, Gustaf Hellsén, Izumi Kaji, James R. Goldenring, Thaher Pelaseyed

## Abstract

A dense glycocalyx, composed of the megaDalton-sized membrane mucin MUC17, coats the microvilli in the apical brush border of transporting intestinal epithelial cells, called enterocytes. The establishment of the MUC17-based glycocalyx in the mouse small intestine occurs at the critical suckling-weaning transition. The enterocytic glycocalyx extends 1 µm into the intestinal lumen and prevents the gut bacteria from directly attaching to the enterocytes. To date, the mechanism behind apical targeting of MUC17 to the brush border remains unknown. Here, we show that the actin-based motor proteins MYO1B and MYO5B, and the sorting nexin SNX27 regulate the intracellular trafficking of MUC17 in enterocytes. We demonstrate that MUC17 turnover at the brush border is slow and controlled by MYO1B and SNX27. Furthermore, we report that MYO1B regulates MUC17 protein levels in enterocytes, whereas MYO5B specifically governs MUC17 levels at the brush border. Together, our results extend our understanding of the intracellular trafficking of membrane mucins and provide mechanistic insights into how defective trafficking pathways render enterocytes sensitive to bacterial invasion.

## Introduction

The epithelium of the small intestine consists of a tight single layer of highly polarized epithelial cells, covered by a permeable mucus layer [1, 2]. Luminal bacteria that breach the mucus layer encounter a second line of defense in a 1 µm thick glycocalyx attached to the microvillus-studded apical brush border of transporting intestinal epithelial cells, called enterocytes [3]. Our previous studies identified membrane mucin MUC17 as a major component of the enterocytic glycocalyx [4], which forms a physical barrier that prevents direct bacterial contact with enterocytes [5]. MUC17 is 4493 amino acids long protein with an extracellular PTS-rich domain consisting of recurring proline, threonine, and serine residues organized in 60 tandem repeats [6]. The serine and threonine residues undergo O-linked glycosylation, resulting in an O-glycosylated mucin domain comprising 80% of the mucin’s molecular weight [7]. The mucin domain connects to the transmembrane domain via an evolutionarily conserved sea urchin sperm protein, enterokinase and agrin (SEA) domain that is auto-catalytically cleaved during mucin biosynthesis and serves as a mechanosensor at the cell surface [8–10]. The cytoplasmic tail domain of MUC17 harbors a class I PSD95, DLG1, ZO-1 (PDZ)-binding motif (PBM) [6], which helps PDZ-containing protein 1, PDZK1 to retain MUC17 in the apical membrane. In addition, MUC17 holds two phosphorylation sites with undefined functions [11].

The MUC17-based glycocalyx is replenished every 12-24 hours [12], which is considerably faster than the turnover of individual enterocytes (3-5 days) [13–15]. As a result, enterocytes must carefully regulate the turnover of MUC17 to guarantee the barrier integrity of the glycocalyx. MUC17 turnover is slower in the ileum of germ-free mice compared to colonized mice, suggesting that the commensal gut microbiota plays a role in the renewal of the glycocalyx [16]. Recycling of membrane proteins in enterocytes is a tightly regulated process since it determines the composition of the apical and basolateral membranes, surface receptor activity, and ion transport. Membrane protein trafficking to and from the cell surface is mediated by myosin motor proteins and Rab GTPases, which coordinate the transport and retention of vesicle-borne membrane proteins within specific endosomal compartments [17]. In addition, sorting nexins in early endosomes mediate rapid recycling of proteins back to the plasma membrane [18]. However, the intracellular trafficking pathway of MUC17 in enterocytes is entirely undefined.

Here, we combined quantitative proteomics, protein-protein interaction assays, CRISPR-Cas9-mediated gene deletion and imaging to demonstrate that the myosin motor proteins MYO1B and MYO5B regulate MUC17 targeting to the apical brush border. Moreover, we identified SNX27 as a novel MUC17 interaction partner.

## Results

### A recombinant MUC17 exhibits correct processing and localization in Caco-2 cells

To identify proteins required for MUC17 trafficking in enterocytes, we turned to the Caco-2 colorectal adenocarcinoma cell line. Unpolarized Caco-2 cells differentiate over a period of 21 days to form a polarized monolayer of columnar epithelial cells with a defined brush border membrane [19]. Importantly, Caco-2 cells express negligible levels of endogenous MUC17 [20, 21], allowing us to constitutively express a recombinant MUC17 with an endogenous signal sequence, an N-terminal 3xFlag tag and a mucin domain consisting of 7 PTS-rich tandem repeats (Figure 1A). MUC17(7TR) localized to the tip of Ezrin- and F-actin-positive microvilli in differentiated Caco-2 cells (Figure 1B-C), thereby recapitulating the position of endogenous MUC17 in human and murine enterocytes [5]. Next, we asked whether Caco-2 cells correctly processed MUC17(7TR) by investigating the presence of mature N- and O-glycans (Figure 1D). Boiling prior to SDS-PAGE dissociates MUC17 into a large N- and a smaller C-terminal subunit that we can detect with fragment-specific antibodies (Figure 1A). The C-terminal subunit separated into two bands on SDS-PAGE; an upper band representing mature EndoH-resistant, PNGaseF-sensitive N-glycans and a lower band representing EndoH-sensitive, ER-resident protein species (Figure 1D, left panel). For O-glycan analysis, we took advantage of StcE, a bacterial metalloprotease with high specificity for O-glycosylated mucin domains [22]. We observed a diffuse 450-kDa band that was digested by StcE, thus representing the fully O-glycosylated N-terminal fragment of MUC17 (Figure 1D, right panel).

**Figure 1.**
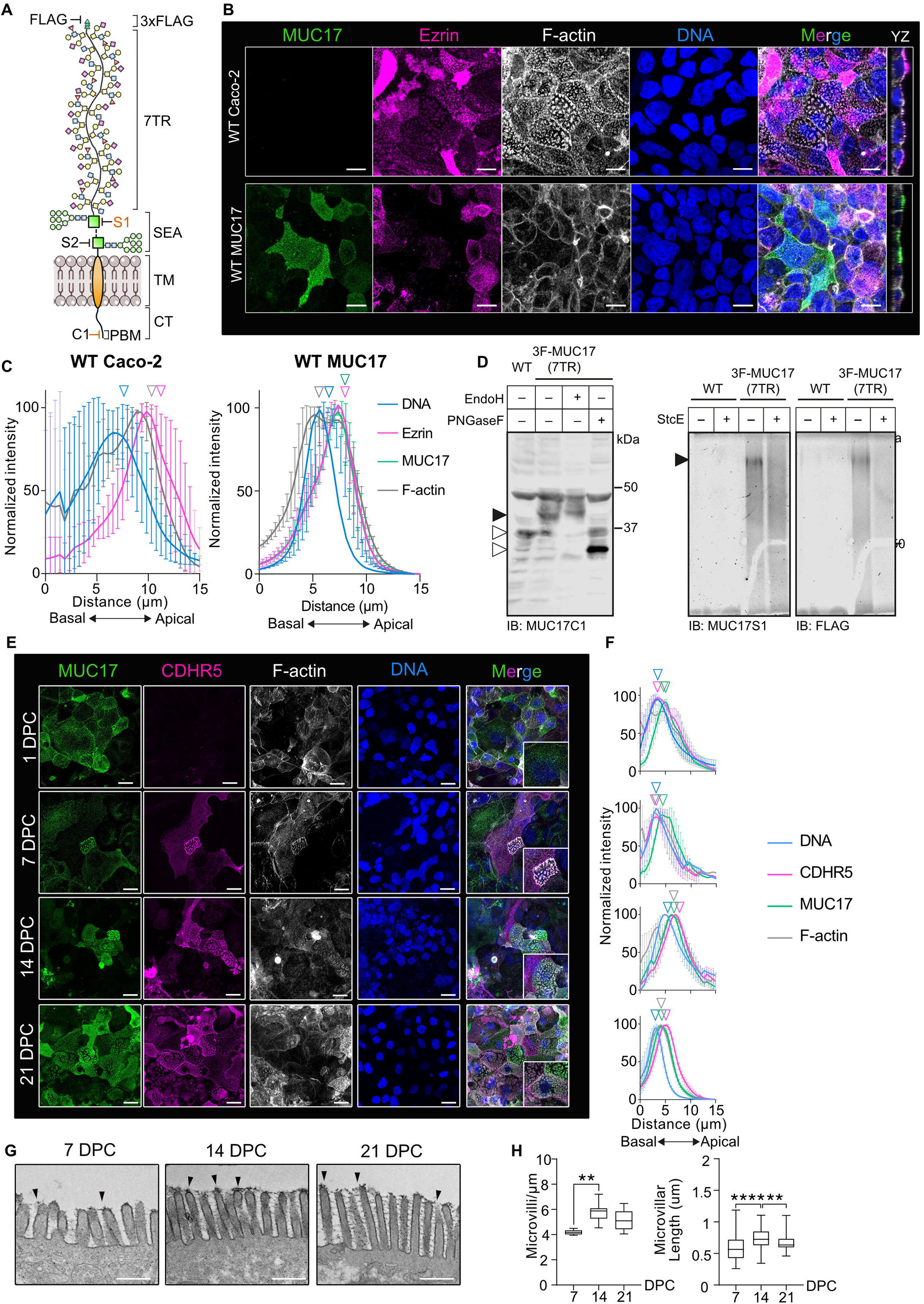
MUC17(7TR) localizes apically in Caco-2 cells. A) Schematic representation of recombinant 3xFlag-MUC17(7TR). TR: tandem repeat, SEA: sea urchin sperm protein, enterokinase and agrin, TM: transmembrane domain, CT: cytoplasmic tail domain, PBM: PDZ-binding motif. Epitopes for the fragment-specific antibodies used in this study are highlighted (FLAG, MUC17-S1, MUC17-S2, and MUC17-C1). Confocal images of Caco-2 cells stained for MUC17, Ezrin, F-actin, and nuclear DNA. Scale bars=10 µm. B) Intensity profiles of MUC17 distribution in WT and MUC17(7TR) Caco-2 cells. n=3 scans and a mean of 73 cells per group. C) Cell lysates of WT and MUC17(7TR) cells treated with PNGaseF, EndoH and StcE, and analyzed by immunoblotting (IB) or in-gel western, and probed with Flag, MUC17-C1, and MUC17-S1 antibodies. Filled and empty arrowheads represent fully mature and immature protein, respectively. D) Assessment of MUC17(7TR) localization during cell differentiation during 1 – 21 dpc. Scale bars=10 µm, n = 3 scans with a mean of 60 cells per time point. E) Intensity profiles of MUC17(7TR) distribution in relation to CDHR5 during cell differentiation. F) TEM micrographs of the glycocalyx during 7, 14 and 21 dpc. Scale bars = 500 µm. G) Microvillar density and microvillar length measured from TEM micrographs. *p< 0.05, ***p<0.001, ****p<0.0001 as determined by one-way ANOVA followed by Tukeýs multiple comparisons test.

Brush border morphology and protein composition change during Caco-2 cell differentiation [23, 24], starting from sparsely spread individual microvilli that culminate in an organized array of tightly packed microvilli, marked by the emergence of an intermicrovillar adhesion complex (IMAC), including CDHR2 and CDHR5, at the tip of microvilli [24]. Since MUC17(7TR) localized to microvillus tips, we asked whether MUC17 localization is influenced by cell differentiation and microvillar packing. Assessment of the localization of MUC17 and CDHR5 during cell differentiation revealed robust apical MUC17 staining independent of cell differentiation stage, whereas CDHR5 appeared after 7 days of cell differentiation (Figure 1E-F). Transmission electron microscopy (TEM) on Caco-2 cells expressing MUC17(7TR) showed that microvillus length and packing, as well as density of the MUC17-based glycocalyx at microvillar tips increased at later stages of cell differentiation (Figure 1G-H). Hence, we concluded that MUC17(7TR) localizes apically to the brush border independent on the differentiation state of Caco-2 cells.

### The interactome of MUC17 uncovers mediators of intracellular trafficking

To identify proteins that participate in trafficking of MUC17, we employed unbiased quantitative proteomics using stable isotope labeling in cell culture (SILAC) and reversible crosslink immunoprecipitation (Re-CLIP) based on nonionic IGEPAL as detergent [25] (Figure 2A, Table S1). Addition of the reversible crosslinker DSP enabled us to capture transient protein interactions. Caco-2 (Light) and Caco-2-MUC17(7TR) (Heavy, C^13^ Lys, C^13^ Arg) cells at 21 dpc, with or without DSP crosslinking, were subjected to Flag-immunoprecipitation and proteomic analysis of the co-precipitated proteins. 35 and 38 confident proteins were identified in the non-crosslinked and crosslinked samples, respectively (Figure 2B-C, Table S1). Out of these proteins, a minority were significantly enriched with MUC17(7TR) and included proteins involved in protein recycling (SNX27), processing (PPM1B, HSPA5, PSMD4, EEF1A1P5, AGR2) and transport (KIF11). PPM1B and AGR2 were only identified in cross-linked samples, whereas SNX27, PSMD4, and EEF1A1P5 were only present in non-crosslinked samples. HSPA5 and KIF11 were found in both conditions (Figure 2D). None of the identified proteins have been reported as MUC17 interaction partners. Due to the small number of identified proteins, we developed a second Re-CLIP protocol based on the ionic detergent SDS to gain a deeper insight into the interactome of MUC17. Using this method, we identified 1058 confident protein hits of which 19 proteins were specifically enriched for MUC17(7TR) (Figure 2E-F, Table S1). Amongst these, we found the myosin motor protein MYO1B, the F-actin bundling protein FSCN1, and proteins associated with the secretory pathway (CKAP4, DPYSL2, HM13, RTN4). Apart from the bait MUC17, we observed differences between the significantly enriched proteins captured by the two Re-CLIP protocols, suggesting that the two methods identify distinct MUC17 interaction profiles (Figure 2G).

**Figure 2.**
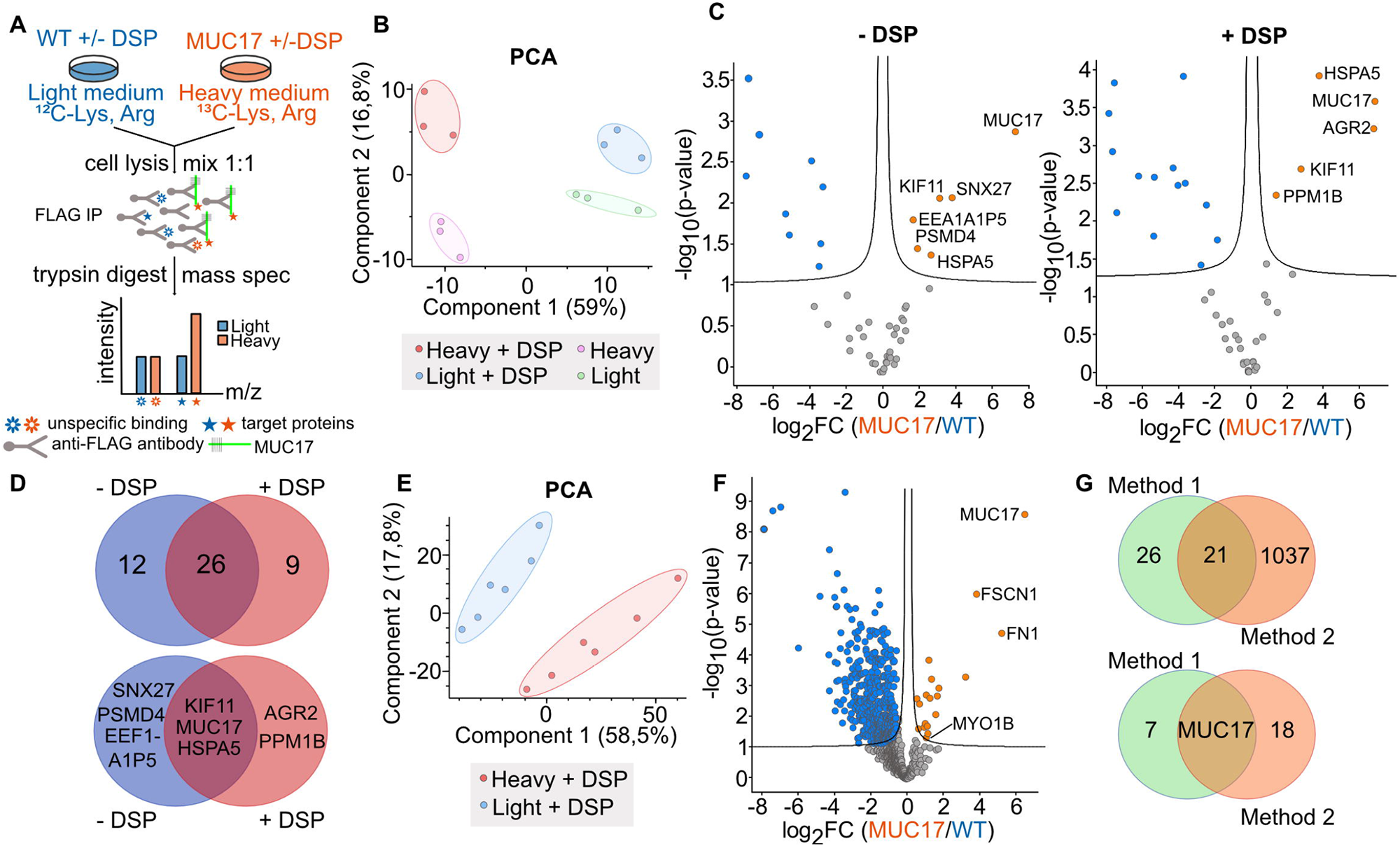
Exploring the interactome of MUC17(7TR) using quantitative proteomics. A) Schematic representation of the workflow for identifying MUC17 interaction partners. B) PCA plot of MUC17(7TR) (Heavy) and WT (Light) samples prepared with IGEPAL (Method 1). C) Volcano plot of MUC17(7TR) (orange) and WT (blue) samples prepared by Method 1. D) Comparisons of all identified proteins (upper) and significantly enriched proteins (lower) identified in C. E) PCA plot of crosslinked MUC17(7TR) (heavy) and WT (light) samples prepared with SDS (Method 2). F) Volcano plot of MUC17(7TR) (orange) and WT (blue) samples prepared by Method 2. G) Comparison of Method 1 and 2 based on all proteins identified (upper) and significantly enriched proteins identified in F.

### MYO1B and SNX27 localize with MUC17(7TR) to the brush border of enterocytes

Based on our interactome discovery, we focused on MYO1B and SNX27. MYO1B regulates the apical targeting of membrane proteins to the brush border [26], whereas SNX27 is involved in the recycling of membrane proteins that interact with its N-terminal PDZ domain [27]. In Caco-2-MUC17(7TR) cells, endogenous MYO1B and SNX27 localized to the apical brush border, where they overlapped with MUC17(7TR) (Figure 3A). At higher magnification, MYO1B localized to the entire length of the microvilli and in the subapical terminal web region, whereas SNX27 localized to microvilli and to distinct puncta within the terminal web region (Figure 3B). Observations in differentiated Caco-2 cells were validated in mouse Ileum, where both MYO1B and SNX27 localized to the brush border where Muc17 is positioned (Figure 3C). For additional validation, we defined the localization of recombinantly tagged rat Myo1b and human SNX27 co-expressed with MUC17(7TR) in differentiated Caco-2 cells (Figure 3D, Figure S1A). HA-Myo1b overlapped with MUC17(7TR) in the apical brush border, whereas SNX27 formed larger puncta below the brush border. MUC17 holds a conserved C-terminal class I PBM that can potentially mediate a PDZ interaction with SNX27 [6, 28]. Therefore, we introduced our tagged constructs into HEK293 cells (Figure S1A) and deployed co-immunoprecipitation assays to investigate whether Myo1b and SNX27 interact with MUC17(7TR). EGFP-SNX27 coprecipitated with MUC17(7TR) (Figure 3E) but, due to the poor expression of EGFP-SNX28ΔPDZ, we were not able to demonstrate if the interaction was mediated by the PDZ domain of SNX27. In conclusion, we demonstrated that both MYO1B and SNX27 localize to the brush border together with MUC17. In addition, we detected a robust interaction between MUC17(7TR) and SNX27, while the MYO1B-MUC17(7TR) interaction was either indirect or too weak to capture without DSP crosslinking (Figure S1B)

**Figure 3.**
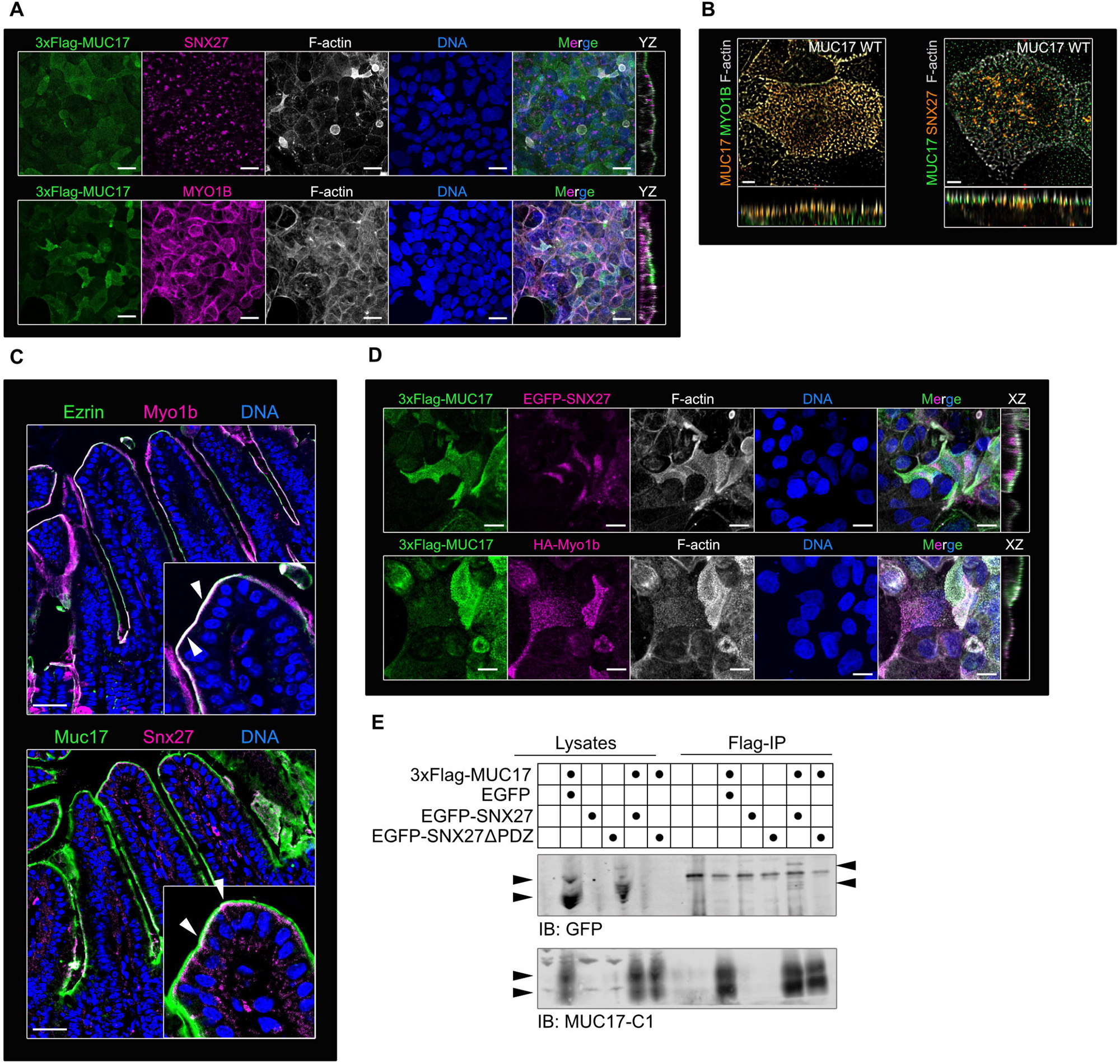
SNX27 interacts with membrane mucin MUC17. A) Confocal images of MUC17(7TR), SNX27 or MYO1B co-stained with F-actin and nuclear DNA. B) High resolution Airyscan images of the brush border of WT MUC17(7TR) cells stained for MUC17(7TR), MYO1B or SNX27, and F-actin. Scale bars = 5 µm. C) Sections of mouse Ileum stained for Ezrin, Myo1b, and nuclear DNA (top) and Muc17, Snx27, and DNA (bottom). Arrows in insets mark the brush border region. Scale bars = 100 µm. D) Confocal images of MUC17 together with recombinant EGFP-SNX27 or HA-Myo1B co-stained for F-actin and nuclear DNA. Scale bars = 5 µm. E) A representative immunoblot of co-immunoprecipitations in HEK 293 cells expressing 3xFlag-MUC17(7TR) and either EGFP-SNX27 or EGFP-SNX27Δ of the total cell lysate whereas 40% of the eluates was loaded on the gel.

### MYO5B regulates MUC17 trafficking to the plasma membrane

Since we discovered the monomeric MYO1B in the interactome of MUC17, we asked whether other non-muscle myosins associate with MUC17. Mining of public single cell RNA-sequencing data sets revealed that MYO1A, MYO5B and MYO7B are highly enriched in enterocytes (Figure S2A). MYO5B is particularly interesting since it transports Rab8^+^Rab11^+^ endosomes carrying membrane proteins to the apical brush border and regulates cell polarity [29, 30]. Moreover, loss-of-function mutations in MYO5B lead to Microvillus inclusion disease (MVID) in humans [31]. To determine the impact of MYO5B on MUC17 trafficking, we stained for Muc17 and Ezrin in ileal sections of *Myo5b^fl/fl^;Vil1-CreERT* mice injected with vehicle or tamoxifen to induce deletion of the *Myo5b* gene in *Vil1*-expressing intestinal epithelial cells. In *Myo5b^fl/fl^;Vil1-CreERT* mice injected with tamoxifen, Muc17 was completely absent from the brush border and restricted to large intracellular vesicles (Figure 4A-B). To further investigate how MYO5B regulates MUC17 transport, we deleted *MYO5B* in Caco-2-MUC17(7TR) cells (Figure S2B-C). While MUC17(7TR) resided in the apical brush border of WT cells, apical MUC17(7TR) was lost in *MYO5B^−/−^* cells, thereby reproducing the phenotype observed in *Myo5b*^Δ^*^IEC^*mice (Figure 4C-E). Moreover, *MYO5B^−/−^*cells demonstrated a dramatic reduction in microvillus-associated Ezrin.

**Figure 4.**
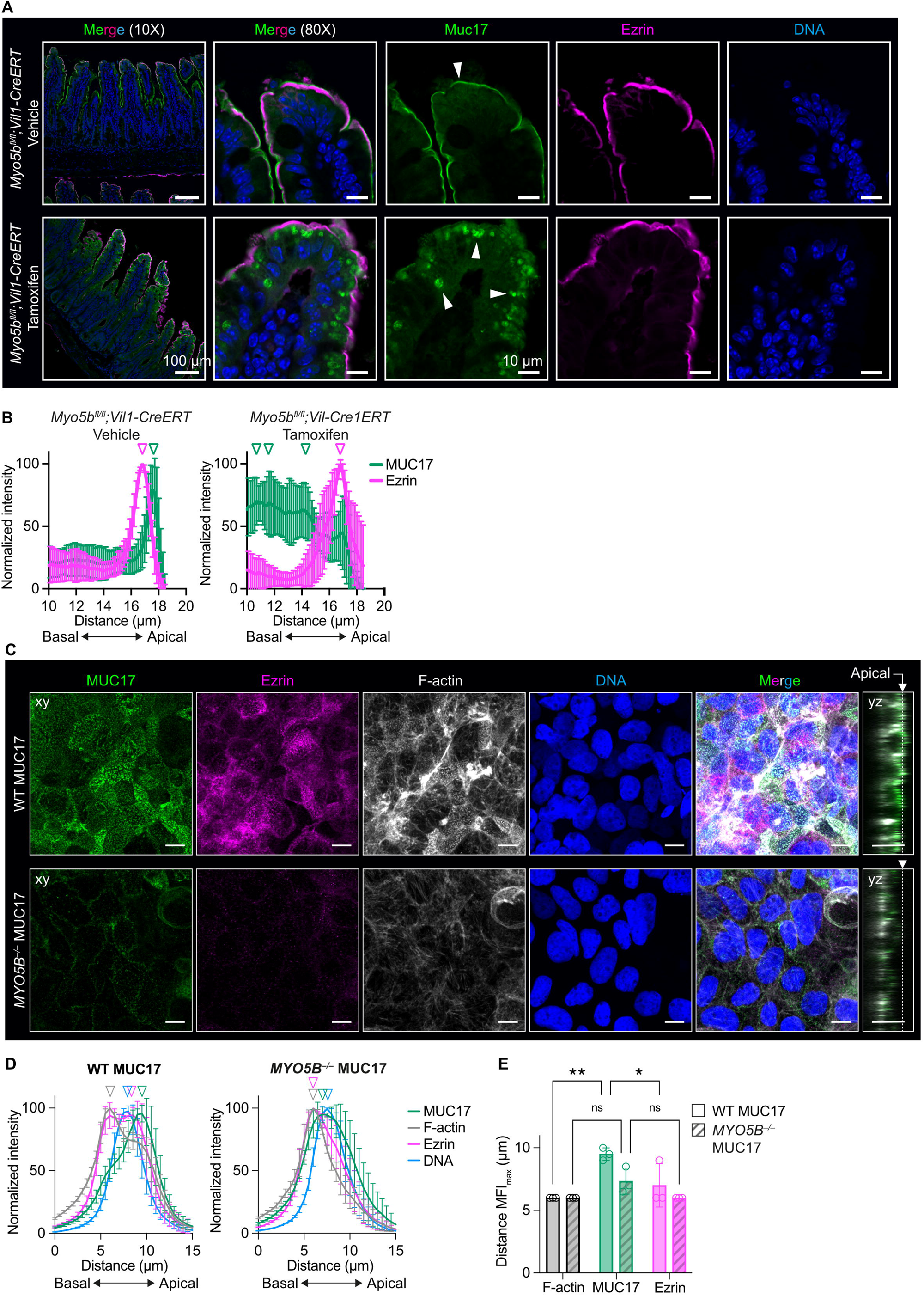
MUC17 resides intracellularly in enterocytes carrying a MYO5B deletion. A) Ileal sections from *Myo5b^fl/fl^;Vil1-CreERT* mice, injected with vehicle or tamoxifen, stained for Muc17, Ezrin and nuclear DNA. B) Intensity profiles of Muc17 and Ezrin in brush border regions in A. C) Confocal images of WT and *MYO5B^−/−^* MUC17(7TR) Caco-2 cells stained for MUC17(7TR), Ezrin, F-actin, and nuclear DNA. Scale bars=20µm. D) Intensity profiles of MUC17(7TR) distribution in WT and *MYO5B^−/−^* Caco-2 cells in relation to Ezrin in C. E) Semi-quantitative analysis of intensity profiles in D. Data are presented as mean ± SD and analyzed by two-way ANOVA corrected for multiple comparison using Sidak. *p<0.05 and **p<0.01.

Due to the dramatic shift in MUC17(7TR) localization observed in *MYO5B^−/−^* cells, we further investigated whether loss of MYO5B affects the surface pool of MUC17(7TR) by applying biotin surface labeling followed by streptavidin affinity purification (Figure 5A). *MYO5B^−/−^* cells presented less MUC17(7TR) on the apical surface compared to the WT control cells (Figure 5B). Next, we took advantage of biotin proximity labelling by antibody recognition coupled to quantitative mass spectrometry to obtain a comparison of the intracellular context surrounding MUC17(7TR) in WT and *MYO5B^−/−^* cells (Figure 5C-E, Table S2). The proximal proteome of MUC17(7TR) in WT cells provided a unique insight into the protein environment that MUC17 encounters during intracellular trafficking to the brush border. The identified proteins participate in brush border and actin cytoskeleton remodeling (MYH14, PLS1, EPS8L2, GSN) and intracellular vesicle trafficking machinery (ANX4, PACSIN3, APPL2, STX3) (Figure 5F, upper panel and 5G). STX3 is particularly interesting since it participates in membrane fusion of endosomes transported by MYO5B. In contrast to WT cells, the proximal proteome of MUC17(7TR) in *MYO5B^−/−^* cells was dominated by proteins associated with basolateral membranes, indicating a disruption of polarized transport of MUC17 to the apical bush border (Figure 5F, lower panel and 5G).

**Figure 5.**
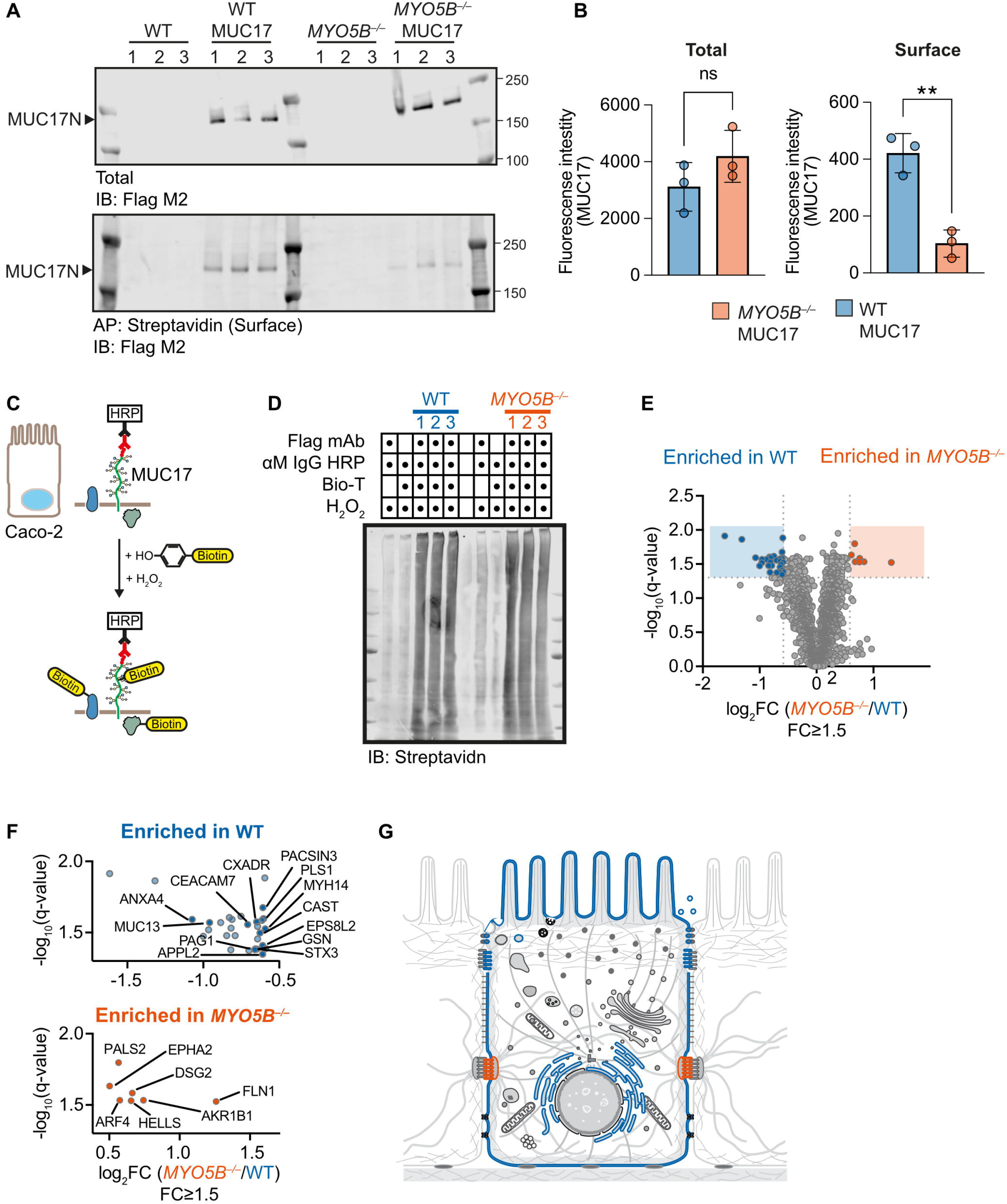
MYO5B regulates MUC17 trafficking to the plasma membrane. A) Surface biotinylation of WT, WT MUC17(7TR), *MYO5B^−/−^* and *MYO5B^−/−^* MUC17(7TR) Caco-2 cells analyzed by immunoblotting. B) Semi-quantitative analysis of band densities in A for total and surface pools of MUC17. Data are presented as mean ± SD. **p<0.01 as determined by unpaired t-test with Welch’s correction, assuming non-equal SD. C) Schematic illustration of biotin proximity labeling of proteins in close proximity to MUC17(7TR). D) Immunoblot of eluates from biotin proximity labeling experiments of WT MUC17(7TR) and *MYO5B^−/−^* MUC17(7TR) Caco-2 cells. E) Volcano plot of proteins enriched in WT MUC17(7TR) and *MYO5B^−/−^* MUC17(7TR) Caco-2 cells from the biotin proximity labeling experiment. F) Significantly enriched proteins for WT MUC17(7TR) (blue) and *MYO5B^−/−^* MUC17(7TR) (orange) Caco-2 cells. G) Visualization of intracellular compartments defined by the proximal proteome of MUC17(7TR) in WT (blue) and *MYO5B^−/−^* (orange) Caco-2 cells.

### MYO1B regulates MUC17 protein levels

Based on the distinct apical and subapical localizations of MYO1B and SNX27, we hypothesized that the two proteins regulate the apical targeting of MUC17. For that reason, we deleted MYO1B and SNX27 separately in Caco-2 cells and re-introduced MUC17(7TR) (Figure S2B, S2D-E). Although MUC17(7TR) remained in the apical brush border in *MYO1B^−/−^* MUC17(7TR) cells (Figure 6A-B), total MUC17(7TR) protein levels were reduced upon loss of MYO1B (Figure 6C). Interestingly, WT MUC17(7TR) cells and *MYO5B^−/−^* cells displayed fewer microvillar clusters compared to WT and *MYO1B^−/−^* cells (Figure 6D). A higher magnification of *MYO1B^−/−^* cells showed that MUC17(7TR) was no longer restricted to the microvilli but also appeared in the terminal web (Figure 6E). Together, our findings demonstrated that deletion of MYO1B resulted in decreased total MUC17 protein levels but did not impact the overall localization of MUC17 in the apical brush border of Caco-2 cells.

**Figure 6.**
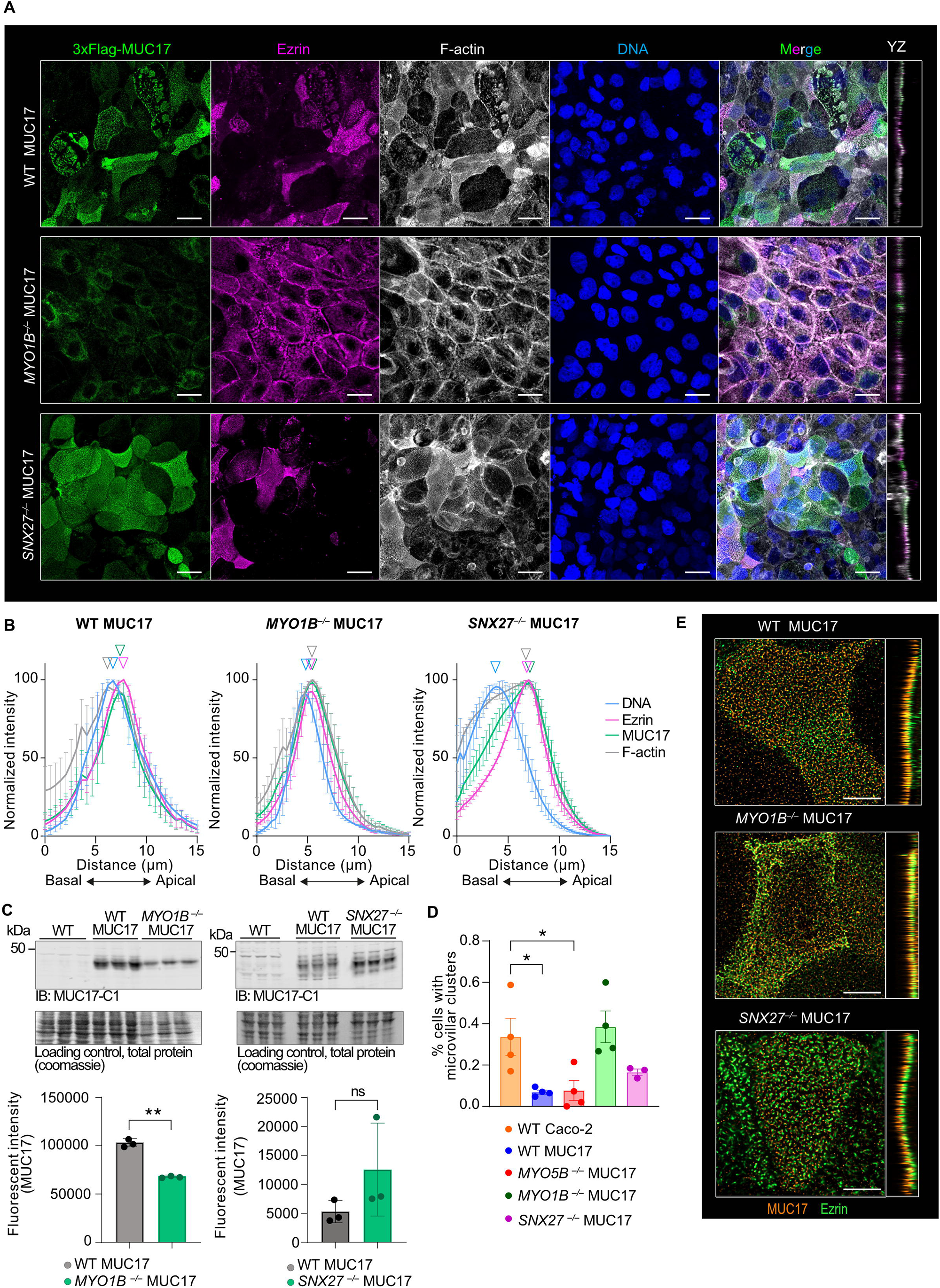
MYO1B regulates MUC17 protein levels. A) Confocal images of WT MUC17(7TR) and *MYO1B^−/−^* MUC17(7TR), and *SNX27^−/−^* MUC17(7TR) Caco-2 cells stained for MUC17(7TR), Ezrin, F-actin, and DNA. Scale bars=20 µm. B) Intensity profiles of MUC17(7TR) distribution in WT and KO cells. n=3 scans and 76 cells per cell line. C) Determination of MUC17(7TR) protein levels in WT, WT MUC17(7TR) and cells lacking either MYO1B or SNX27 (upper panel). Lower panel represents quantitative analysis of protein levels in WT and KO cells. Data are presented as mean ± SD. **p<0.01 as determined by unpaired t-test with Welch’s correction, assuming non-equal SD. D) Quantification of cells displaying microvillar clusters in WT, WT MUC17(7TR) and *MYO5B^−/−^*, *MYO1B^−/−^*, *SNX27^−/−^* MUC17(7TR) cells given as percentage. *p<0.05 as determined by one-way ANOVA and Dunnett’s multiple comparisons test. n = 3-4 scans with a mean of 358 cells measured per cell line. E) High resolution Airyscan images of MUC17(7TR) distribution in relation to Ezrin in the brush border region of WT and *MYO1B^−/−,^ SNX27^−/−^* MUC17 cells. Scale bars = 5 µm.

### Reduced apical MUC17 targeting in MYO1B-deficient cells

While loss of MYO5B had a dramatic negative effect on the targeting MUC17 to the apical brush border, MYO1B and SNX27 had modest effects on MUC17 localization in fixed differentiated Caco-2 cell. Therefore, we asked whether MYO1B and SNX27 influence the kinetics of MUC17 at the apical brush border in live cells. To address this question, we took advantage of Fluorescence recovery after photobleaching (FRAP). To specifically tag surface-attached MUC17(7TR) with a fluorescent label, we coupled a fluorescent dye to the inactive E447D mutant of StcE that only binds mucins. First, we showed that StcE E447D bound and enriched for the mature and fully glycosylated form of MUC17(7TR) with a molecular weight of around 450 kDa in WT cells (Figure 7A-B, Figure S3). StcE E447D also captured mature MUC17(7TR) in *MYO5B^−/−^* and *SNX27^−/−^* cells, and to a lower extent in *MYO1B^−/−^* since total MUC17(7TR) protein levels are lower in this line (Figure 6C). Thus, we concluded that the difference in MUC17 protein levels and localization in KO cells was not a result of altered MUC17(7TR) processing. Next, we used imaging to investigate if fluorescently labelled StcE E447D specifically detects surface-bound MUC17(7TR) in Caco-2 monolayers (Figure 7C). Strikingly, while the staining with StcE E447D in WT Caco-2 appeared to be mainly intracellular, there was a strong correlation between MUC17(7TR) and StcE E447D in the apical brush border of MUC17(7TR) cells (Figure 7C-E). In MUC17(7TR) cells lacking MYO1B, MYO5B or SNX27, there was less overlap between MUC17(7TR) and StcE E447D (Figure 7E). After demonstrating that StcE E447D specifically binds MUC17(7TR), we used FRAP to measure the kinetics of MUC17 at the apical brush border in live WT and KO cells (Figure 7F). Fluorescent recovery of MUC17(7TR) in WT cells was slow (t_1/2_=6 min) and incomplete at the end point of the experiment (12 minutes) (Figure 7F and 7G, left panel). In addition, we estimated the mobile fraction representing the lateral diffusion of MUC17(7TR) within the plasma membrane to 45% (Figure 7G, right panel). Compared to WT cells, the recovery of MUC17(7TR) was significantly slower in *MYO1B^−/−^* cells (t_1/2_=10 min), while MUC17(7TR) recovered faster in *SNX27^−/−^* cells. Moreover, the mobile fraction of MUC17(7TR) decreased by 50% in *SNX27^−/−^* cells. No difference in turnover was observed in MUC17(7TR) *MYO5B^−/−^* cells. Together, our results indicate that MUC17 trafficking and targeting to the plasma membrane is a slow process under baseline conditions with limited diffusion at the brush border. Furthermore, MYO1B regulates apical targeting of MUC17, whereas SNX27 affects the dynamic behavior of MUC17 at the plasma membrane.

**Figure 7.**
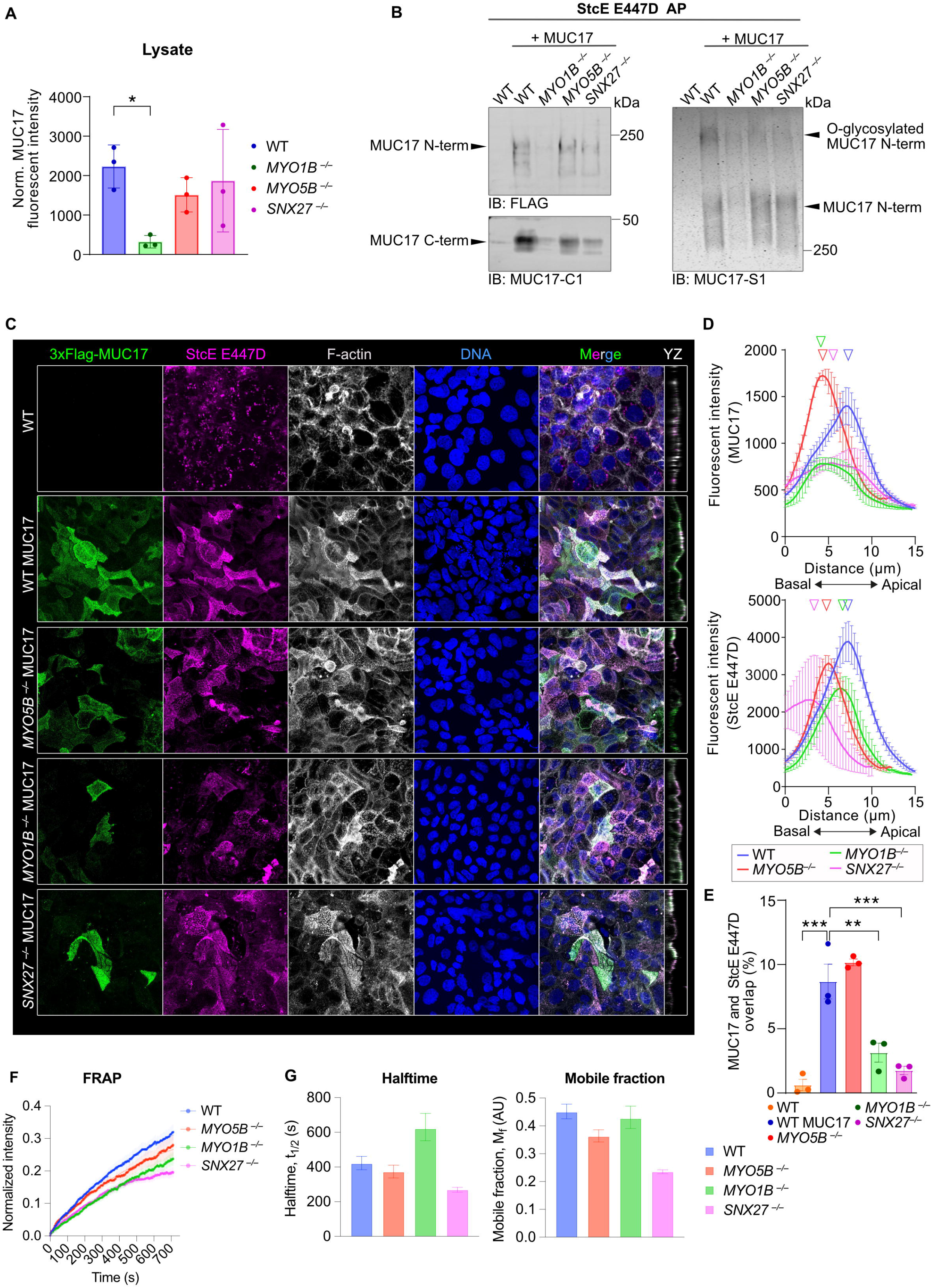
Reduced apical MUC17 targeting in MYO1B-deficient cells. A) Densitometric analysis of total MUC17(7TR) normalized to total loaded protein for each cell lysates used for StcE AP in B. Data are presented as mean ± SD *p<0.05 as determined by Kruskal Wallis and Dunn’s multiple comparisons test. B) Assessment of maturation of MUC17(7TR) by affinity purification using StcE E447D in WT, MUC17(7TR)-expressing WT and KO Caco-2 cells. Eluates (40%) were analyzed by immunoblotting (left and middle panel) or in-gel western (right panel) using Flag, MUC17-C1 or MUC17-S1 antibodies. C) Confocal images of WT, MUC17(7TR) or *MYO5B^−/−,^ MYO1B^−/−,^ SNX27^−/−^* MUC17(7TR) cells stained with MUC17 and fluorescently conjugated StcE E447D, alongside YZ orthogonal projections. D) Intensity profiles of MUC17(7TR) (upper) and StcE E447D (lower) for WT and KO cells in C. n=3 scans and a mean of 82 cells per cell line. E) Proportion of MUC17(7TR) signal that overlaps with StcE E447D in C. Data are presented as mean ± SD and analyzed by two-way ANOVA corrected for multiple comparison using Sidak, *p<0.05, **p<0.01, ***p<0.001. F) Curves representing recovery after photobleaching of MUC17(7TR)-bound StcE E447D in the plasma membrane. n=19-23 per group. G) Halftime and Mobile fraction for WT and KO cells extracted from F. Data for each group are represented with 95% confidence intervals.

## Discussion

We have previously identified a MUC17-based glycocalyx covering the surface of enterocytes. Importantly, the absence of the enterocytic glycocalyx results in increased bacterial contact with the enterocytic brush border [5]. Our findings highlight the importance of a precise regulation of MUC17 trafficking in enterocytes. In this study, we mapped the trafficking machinery of MUC17 in the epithelial cell line Caco-2. Confluent Caco-2 cells undergo a differentiation process, mirroring the *in vivo* maturation of enterocytes along the crypt-villus axis. Differentiated Caco-2 cells are characterized by a less pronounced tumorigenic phenotype, cytoskeletal rearrangements [23] and tightly packed microvilli held together by tip links formed by an IMAC [24]. Our investigation showed that MUC17 is expressed on the apical membrane regardless of the cell differentiation whereas the MUC17-based glycocalyx becomes denser as microvillus packing takes place late during cell differentiation. These findings are consistent with the localization of MUC17 in differentiated villus enterocytes as well as less differentiated crypt cells in human and mouse ileum [5]. These sequences of events suggest that MUC17 is inserted in the apical membrane prior to microvillus assembly. As the enterocyte reaches a differentiated state marked by a brush border, tight packaging of microvilli places MUC17 at the distal tip of individual protrusions, where a dense glycocalyx is established.

PDZ proteins regulate membrane mucin trafficking through the Golgi apparatus and retention at the apical cell membrane [6, 32] but we lack a comprehensive understanding of how large membrane mucins such as MUC17 are transported within enterocytes. Here, we developed two enrichment protocols that both identified primarily cytoplasmic proteins involved in protein recycling, processing, and transport. PDZ-domain containing SNX27 directs endocytosed proteins from the early endosomes to the plasma membrane [33]. We showed that MUC17 binds SNX27 and that both proteins localize to the apical brush border of enterocytes, but we could not confirm that the interaction was PDZ-dependent. We also showed that SNX27 impacts the slow kinetics of MUC17 at the apical brush border. In *SNX27^−/−^* cells, a higher proportion of MUC17 remained static within the brush border, while the remaining mobile fraction diffused more rapidly. Our data indicate that enterocytes maintain glycocalyx barrier integrity by stabilizing the apical pool of MUC17 in the absence of SNX27-mediated recycling of membrane mucins to the plasma membrane.

The role of the unconventional non-muscle myosins in microvillar assembly and function has been extensively characterized [34]. Our mapping of the MUC17 interactome identified the monomeric myosin MYO1B, which regulates the targeting of amino acid transporters to the brush border of kidney cells [26]. MYO1B interacts with actin filaments but lacks the capacity to transport vesicular cargo along filaments [35, 36]. Actin-bound MYO1B induces tubule formation in endosomal and lysosomal membranes, as well as the trans-Golgi network, thereby controlling proteins trafficking between endocytic compartments [37, 38]. Both endogenous and recombinant MYO1B localized with MUC17 in the brush border of cultured cells and mouse Ileum. Depletion of MYO1B resulted in reduced total MUC17 protein levels and slower MUC17 turnover in the brush border. The latter phenotype could be explained by a higher abundance of microvillar clusters that restrict MUC17 diffusion in the brush border membrane of *MYO1B^−/−^* cells. We also addressed the role of the dimeric unconventional myosin MYO5B in MUC17 trafficking. Apical transporters such as NHE3, AQP7, and SGLT1 depend on MYO5B for their targeting to the brush border [39] and mutations in the *MYO5B* gene have been linked to the congenital diarrheal disorder MVID caused by the mislocalization of membrane transporters that maintain cell and fluid homeostasis [40]. MUC17 was removed from the enterocytic brush border in the ileum of *Myo5b*^Δ^*^IEC^*mice and in *MYO5B^−/−^* Caco-2 monolayers. Moreover, biotin proximity labelling in *MYO5B^−/−^*cells revealed that the recycling defects associated with MYO5B deletion resulted in incorrect targeting of MUC17 to the basolateral membrane domain. Our findings are in line with the role of MYO5B in regulating apical cell polarity [41].

In conclusion, we show that MYO1B, MYO5B and SNX27 regulate the intracellular trafficking of the glycocalyx-forming membrane mucin MUC17 in enterocytes. Unravelling the cellular mechanisms that govern the formation of the glycocalyx barrier sheds light on fundamental cellular processes for combatting bacterial encroachment on the intestinal epithelium.

Importantly, our insights into MUC17 trafficking pathways could prove critical for identifying molecular defects that render enterocytes sensitive to bacterial invasion.

## Material and Methods

### Antibodies and fluorescent probes

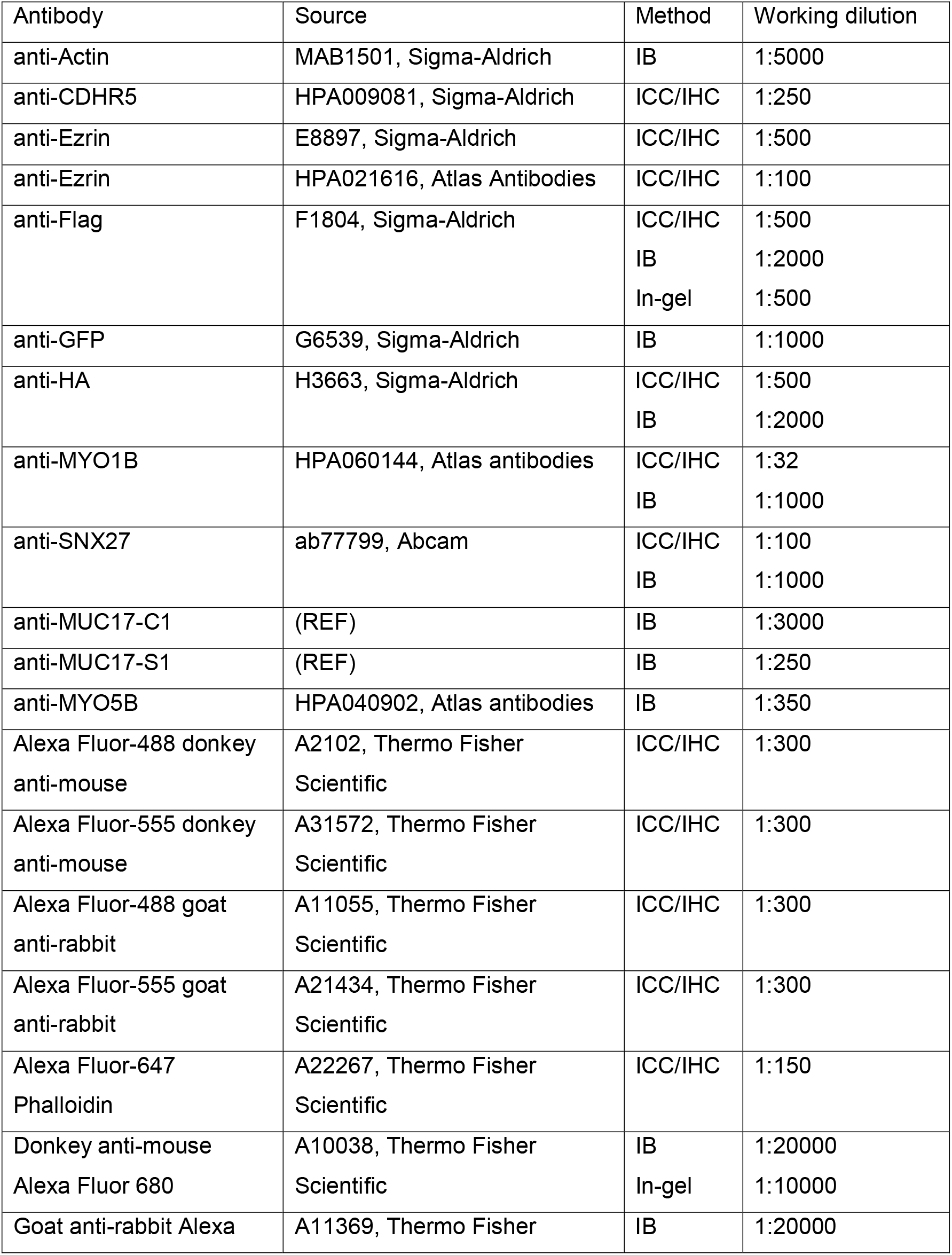
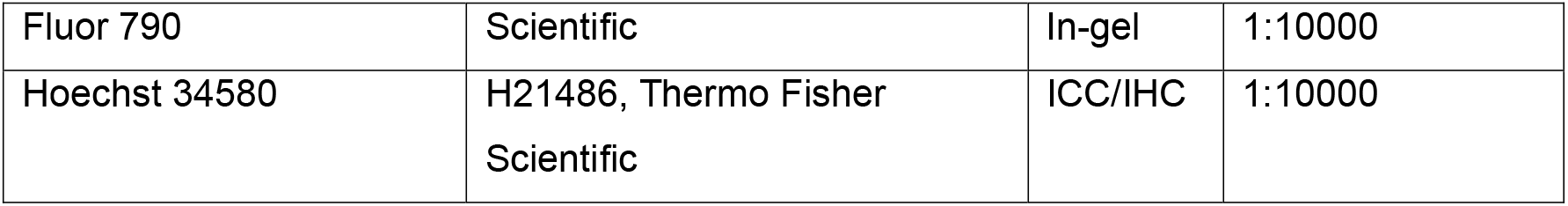

### Plasmids

cDNA encoding recombinant MUC17(7TR) with N-terminal 3xFlag tag was generated using Gibson Assembly (E2611S, NEB) following the manufactures protocol. A cDNA insert with the endogenous MUC17 signal sequence fused to 3xFlag followed by sequence-optimized last 7 N-terminal tandem repeats of the mucin was prepared (see Supplementary figure S4). The insert was equipped with 5’ and 3’ flanking sequences overlapping with the pXL-CAG-Zeocin-3xF2A plasmid [42] digested with NotI and AscI restriction enzymes. Rat Myo1b (Plasmid #135064, Addgene) C-terminal Myc-tag, and SNX27 and SNX27ΔPDZ (a kind gift by Prof. Peter J. Cullen [28, 33]) with N-terminal EGF-tag were cloned into the pXL-CAG-Zeocin-3xF2A. The following guide RNAs were used for the deletion of *MYO1B*, *MYO5B* and *SNX27* genes using the pLentiCRISPR v2 vector according to the protocol by the Zhang lab [43, 44] *MYO1B* 5’-ATGAAGGTCTCCTCATTGAG-3’, *MYO5B* 5’-GCGCTCAGCTGAGTTAACCA-3’, and *SNX27* 5’-GCTACGGCTTCAACGTGCG-3’. Vectors containing gRNAs were transformed into One Shot™ Stbl3™ Chemically Competent E. coli according to manufactureŕs protocol (Invitrogen) and confirmed by sequencing using primer U6-F GAGGGCCTATTTCCCATGATT.

### Immunohistological sections from human and mouse ileum

Human biopsies were sampled from ileum of individuals without suspected Inflammatory Bowel Disease, who were referred to Sahlgrenska University Hospital (Gothenburg, Sweden) for ileocolonoscopy, and subject to the provision of written informed consent (ethical permit 2020-03196). Wild-type C57BL/6N mice were maintained under standardized conditions of temperature (21–22°C) and illumination (12-hour light/dark cycle) with food and water ad libitum. The Swedish Laboratory Animal Ethical Committee in Gothenburg approved the experiments conducted in this study (ethical permit 2285-19). The care and use of animals were performed in accordance with the Swedish animal welfare legislation, which meets the European Convention for the Protection of Vertebrate Animals used for Experimental and other Scientific Purposes (Council of Europe N° 123, Strasbourg 1985) and the European Union Directive 2010/63/EU on the protection of animals used for scientific purposes. Animals were anesthetized with isoflurane followed by cervical dislocation. Animals of 6-8 weeks of age and of both genders were used. Weaning occurred on day 21 after birth (P21). For investigation of Myo5b function in mouse ileum, Cre recombinase was activated in 8-to 10-week-old *VillinCreErt2;Myo5b^fl/fl^* by one intraperitoneal injection of tamoxifen (2 mg).

Tamoxifen-injected *Myo5b^fl/fl^* mice and *VillinCreErt2;Myo5b^fl/+l^* mice were used as controls [29]. All *VillinCreErt2;Myo5b^fl/fl^* and control mice were killed 4 days after the tamoxifen dose. The care, maintenance, and treatment of *VillinCreErt2;Myo5b^fl/fl^* mice followed protocols approved by the Institutional Animal Care and Use Committee of Vanderbilt University.

### Cell culture, transfections and Crispr/Cas9-mediated gene deletion

Caco-2 (ATCC HT-37) and HEK293T (ATCC CRL-1573) cells were cultured in Iscovés modified Dulbeccós medium (IMDM, Invitrogen Life Technologies, Carlsbad CA) containing 10% (vol/vol) FBS at 37°C in 5% CO2. Caco-2 cells for SILAC (#A33969, #88210 Thermo Fisher Scientific) were cultured in Dulbeccós modified Eagle medium supplemented with 13C6 L-lysine and ^13^C_6_L-Arginine (heavy medium) or ^12^C_6_L-Lysine and ^12^C_6_L-Arginine (light medium) respectively for 5 passages to ensure complete incorporation (>95%). Transfections to generate stable clones were performed with 50% and 80% confluent Caco-2 and HEK293T cells, respectively, seeded in 9.6 mm^2^ wells. All transfections to introduce recombinant constructs were performed using Lipofectamine® 2000 to introduce the PiggyBac transposon system with a transposase (pCAG-mPB-orf):transposon (pXL-BacII-CAG-Zeocin-triple-F2A) ratio of 1:2.5. Transfected cells were incubated with a total of 4 µg DNA and 10 µL Lipofectamine 2000 (11668019, Thermo Fisher Scientific) complex for 72 hours and selected for another 14 days with 300µg/mL Zeocin. For CRISPR/Cas9-transfections, cells were selected for 3 days with 700 µg/mL G418, and individual colonies picked for expansion and screening by immunoblotting.

### Co-immunoprecipitation

#### Method 1 with nonionic IGEPAL detergent

3 × 10^5^ of MUC17(7TR) Caco-2 (Heavy) and WT Caco-2 (Light) cells were seeded into 9.6 mm^2^ wells and used for pull-downs experiments at 14 DPC. Re-CLIP was performed by rinsing the cells three times with 37°C PBS followed by incubating cells with 1.25 mM DSP for 2 min at 37°C. DSP was quenched at RT with 3 washes of TBS. Non-crosslinked cells were washed three times in TBS at RT. Cells were subjected to Flag-immunoprecipitations as previously described [45]. Briefly, cells were washed 3x5 min in 37 °C PBS, lysed in 1.5 µL ice-cold Lysis buffer 1 (0.5 % IGEPAL, 250 mM NaCl, 50 mM Tris/HCl pH 7.4, 1 mM EDTA, 1X cOmplete EDTA-free protease inhibitor cocktail (11697498001, Roche), 1mM PMSF, 15 µL phosphatase inhibitor cocktail 2 (P5726, Sigma-Aldrich) and 3 (P0044, Sigma-Aldrich)), and incubated at 4°C for 20 min on an orbital shaker. Cells were collected by scraping and sonicated for 3 min in a water-bath sonicator with ice at 10°C. The cell lysate was cleared by centrifugation at maximum speed for 30 min at 4°C and 50 µL of the supernatant was saved as input. 50 µL of EZview ™ Red ANTI-FLAG® M2 affinity gel (F2426, Sigma-Aldrich), equilibrated in Lysis buffer 1, was added to the sample that were incubated overnight at 4°C on rotation. Beads were washed three times in Lysis buffer 1 and three times in Lysis buffer 1 without IGEPAL at 4°C. Enriched proteins were eluted by adding 25 µL 3xFLAG peptide (4 µg/µL) for 30 min at 4°C. Eluted proteins were separated from beads with Corning Costar Spin-X filter units (#CLS8162, Sigma-Aldrich) and stored at -20°C until further processing.

#### Method 2 with ionic SDS detergent

Cells were crosslinked with 1.25 mM DSP for 2 min at 37 °C followed by three washes in TBS. Cells were lysed in 500 µL Lysis buffer 2 (1% SDS, 250 mM NaCl, 50 mM Tris/HCl pH 7.4, 1mM EDTA, 1x cOmplete EDTA-free protease inhibitor cocktail, 1mM PMSF, 3 µM Beta-Glycerol phosphate, 15 µL phosphatase inhibitor cocktail 2 and 3). Cell lysates were sonicated for 10 seconds and centrifuged at maximum speed for 30 min at 4°C. Supernatants from heavy and light samples were mixed at a 1:1 ratio and diluted 1:10 to reach 0.1% SDS final concentration. 50 µL EZview ™ Red ANTI-FLAG® M2 affinity gel was added to each sample and incubated overnight at 4°C on rotation. Beads were washed three times in TBS+0.1% SDS and eluted by the addition of 50 µL elution buffer (1% SDS, 100 mM Tris pH 8.0, 100mM DTT) and boiling at 95°C for 5 min. Eluted proteins were separated from beads with Costar Spin-X filter units (8160, Corning) and stored at -20°C until further processing.

### Co-immunoprecipitations in HEK293 cells

6 × 10^5^ HEK293T cells stably expressing recombinant constructs were seeded in 9.6 mm^2^ wells and used for pull-down experiments. Co-immunoprecipitations were performed in the absence of DSP according to method 1 with the additional step of blocking the EZview ™ Red ANTI-FLAG® M2 affinity gel with 5% BSA in TBS for 2 hours at 4°C on rotation and washed 2 times in Lysis buffer 1 without IGEPAL before performing immunoprecipitation.

### Cell surface biotinylation

3 × 10^5^ Caco-2 cells were seeded on 9.6 mm^2^ wells and let to differentiate for 14 days post-confluency (DPC). To label the surface proteins, cells were washed three times with ice-cold PBS and incubated with 0.25 µg/mL biotin hydrazide (66640-86-6, Sigma-Aldrich) in cold PBS for 1 hour on ice. After biotin labeling, cells were washed three times with TBS (pH 7.4) and left in the last TBS wash for 20 min at RT. Cells were lysed in 1 mL Lysis Buffer (1% Triton-X, 25 mM Tris/HCl pH 7.4, 150 mM NaCl, 1 mM EDTA, 4% glycerol supplemented with 1X cOmplete EDTA-free protease inhibitor cocktail) for 10 min on ice. Lysates were homogenized by sonication and centrifuged at 16,000 g for 30 min at 4°C. 50 µL of cell lysate was mixed with reducing sample buffer and used as input loading control. 30 µL EZview™ Red Streptavidin Affinity Gel (E5529, Sigma-Aldrich) was added to each supernatant and incubated on rotation for 2 hours at 4°C. After three washes with Lysis buffer, the bound material was eluted with reducing sample buffer and boiled at 95°C for 5 min. Samples were separated on a 4–15% gel by SDS-PAGE and transferred to PVDF-FL membranes (IPFL00010, Merck). Membranes were blocked with 5% non-fat milk in PBS and incubated with primary antibodies diluted in 5% non-fat milk in PBS+0.1%Tween-20 (PBS-T) O/N at 4°C. After 3 PBS-T washes, protein bands were visualized with Odyssey CLx imaging system (LI-COR Biosciences). Total biotinylated proteins in samples were detected with Alexa Fluor™ 790 Streptavidin Conjugate (1:20 000, S11378, Thermo Fisher Scientific). Band densities were quantified using Image Studio quantification software (LI-COR Biosciences).

### Expression and labelling of StcE and StcE E447D

Tuner (DE3) competent cells (70623, Sigma-Aldrich) were transformed with pET28b-StcE-Δ35-NHis or pET28b-StcE-E447D-Δ35-NHis (kind gift from Prof. Carolyn Bertozzi [46]) and grown on LB Agar with kanamycin at 37 °C overnight. A single colony was pre-cultured in 10 mL LB kanamycin overnight, and the preculture expanded in 1 L LB kanamycin until an optical density of 0.85 was reached. Protein production was induced with 0.2 mM IPTG at 30°C overnight. Bacterial cells were centrifuged at 3500 × g for 20 min at 4°C, resuspended in 20 mL of ice-cold PBS, and centrifuged at 3500 × g for 20 min at 4°C. Cell pellets were resuspended in 20 mL of ice-cold Binding Buffer (20 mM sodium phosphate, 300 mM NaCl, 20 mM Imidazole, pH 7.4.) containing 2.5X Roche Complete EDTA-free protease inhibitor cocktail (11873580001, Sigma-Aldrich). The bacterial slurry was sonicated for 8 × 30 seconds at 50% duty in a water bath maintained at 4°C. Lysates were centrifuged at 22,000 × g at 4°C for 20 minutes and poured over 4 mL of HisPur Cobolt resin (89964, Thermo Fisher Scientific). The slurry was rotated at 4°C for 1 hour and spun down at 700 × g for 2 minutes. The resin was washed three times with Binding Buffer including protease inhibitor cocktail, and bound protein eluted at 4°C with three subsequent 15-minute elutions using 3 mL of Elution Buffer (20 mM sodium phosphate, 300 mM NaCl, 500 mM Imidazole, pH 7.4). Elution fractions were pooled and dialyzed against 5 L PBS at 4°C overnight, followed by a second round of dialysis in 5 L PBS for 4 hr at 4°C.

For enrichment of MUC17(7TR) using StcE E447D, StcE E447D was couped to CNBr-Activated Sepharose 4B (17043001, Cytiva). 1 gram of CNBr-Activated Sepharose 4B was resuspend in 10 mL of 1 mM HCl for 1 hour, centrifuged at 1000 × g for 5 min at RT and washed for 15 minutes with 10 mL of 1 mM HCl followed by centrifugation. Swelled agarose was washed with 2 × 10 mL Coupling Buffer (100 mM NaHCO_3_, 500 mM NaCl, pH 8.3) and centrifuged between each wash. 10 mg of StcE E447D was diluted with Coupling Buffer to a final volume of 10 mL, added to the swelled CnBr agarose and rotated overnight at 4°C. The CnBr agarose was washed with 2x 10 mL Coupling Buffer and quenched with 10 mL ice-cold 250 mM Glycine on overnight rotation at 4°C. The CnBr agarose was washed 5 times with 10 mL Coupling Buffer and resuspended in 10 mL H buffer (150 mM NaCl, 50 mM Tris pH 7.4 + 0.02% NaN_3_). The volume was adjusted to a 50% slurry. 50-100 µl of 50% StcE-CnBr slurry was used for each pull-down.

### EndoH, PNGaseF and StcE treatments

3 x10^5^ Caco-2 cells were seeded in 9.6 mm^2^ dishes and differentiated for 14 DPC. Cells washed 3 × 5 min with PBS (RT) and lysed with 200 µL ice cold Lysis buffer (25 mM Tris-HCL pH 7.4, 150 mM NaCl, 4% glycerol, 1% Triton X-100) complemented with final concentration of 1X EDTA-free Complete protease inhibitor cocktail (34044100, Roche) and 1 mM PMSF (78830, SigmaAldrich). Cells were incubated with Lysis buffer for 10 min on ice, collected by scraping and homogenized by sonication in water bath at 40% amplitude for 30 sec pulses for 4 min. Cell lysates were cleared by centrifugation at 16,000 × g for 30 min at 4°C. For EndoH and PNGaseF treatments, 30 µL cell lysates were mixed with 10µL 200 mM DTT, 1 µL PMSF, 2 µL 3.0M NaAc pH 5.4 (only for EndoH treatment), 5 µL EndoH (11643053001, Sigma) or 6 µL PNGaseF (11365177001, Sigma) and diluted with Lysis buffer to 50 µL. Untreated control samples were prepared without the addition of NaAc, EndoH or PNGaseF. For StcE treatment, 42 µL cell lysate was mixed with 1 µL active StcE (5.8 mg/mL). All samples were incubated at 37°C overnight and reduced in 4X reducing sample buffer (8% SDS, 400mM Dithiothreitol, 40% glycerol, 200 mM Tris pH 6.8, 0.4% bromophenol blue) followed by boiling at 95°C for 5 min.

### Immunoblots and in-gel western blots

Samples were separated on precast 4%–12% SDS polyacrylamide gel (XP04125BOX, ThermoFisher Scientific). Proteins were transferred to a PVDF-FL membrane (05317, Millipore) with a current of 2.5 mA/cm^2^ for 1 h. Membrane was blocked in 5% non-fat milk in PBS for 30 min and incubated with primary antibodies diluted in 5% non-fat milk in PBS + 0.1% Tween-20 (PBS-T) overnight at 4°C. Membrane was washed three time in PBS-T and incubated with secondary antibodies diluted in 5% non-fat milk in PBS-t + 0.02% SDS for 1 hour at RT in the dark. Membrane was washed three time in PBS-T and visualized on an Odyssey CLx near infrared fluorescence imaging system (LI-COR Biosciences). Protein quantification was performed using Image Studio quantification software (LI-COR Biosciences). For Coomassie stains, membranes were stained with Imperial Protein Stain (24615, Thermo Scientific), destained in 5% MeOH and 7% Acetic Acid, and visualized on an Odyssey CLx near infrared fluorescence imaging system (LI-COR Biosciences).

Samples for In-gel westerns were separated on precast 4%–12% SDS polyacrylamide gel. Proteins were fixed by 50% isopropanol + 5% acetic acid in ultrapure water for 15 min. Gel was washed extensively in ultrapure water 3x15 min and incubated with primary antibodies diluted in 5% BSA in PBS overnight at 4°C. After 3x10 min washes in PBS-T, gel was incubated with secondary antibodies diluted in 5% BSA in PBS+0.1% Tween^®^ 20 for 2 hours (RT). Gel was washed 3x10 min in PBS-T and visualized on an Odyssey CLx near infrared fluorescence imaging system.

### Biotin proximity labeling by antibody recognition

3 × 10^5^ Caco-2 cells seeded on 9.6 mm^2^ wells and differentiated for 14 days were washed 2 times with PBS. Cells were fixed with 4% paraformaldehyde in PBS for 15 min at RT and washed twice in PBS-T. Next, cells were permeabilized in PBS + 0.5% Triton X-100 for 7 min at RT followed by 3 × 10 min washes with PBS-T. 30 mM H_2_O_2_ in PBS was added overnight at RT to quench endogenous peroxidase activity. Another 30 mM fresh H_2_O_2_ in PBS was added for 10 min followed by two 10 min washes with PBS-T. Cells were incubated with blocking buffer (5% BSA in PBS) for 2 hours on a shaker and stained with Flag mAb 1:500 diluted in blocking buffer overnight at 4°C in a humid chamber on an orbital shaker. After three subsequent 1-hour PBS-T washes, cells were incubated with 1:1000 goat anti-mouse HRP diluted in blocking buffer for 1 hour at RT. Unbound antibody was removed by three 2 hours washes with PBS-T and cells pre-incubated with 500 μ concentration) at RT. After 10 min, a final concentration of 2.5 mM H_2_O_2_ in PBS was added to the Biotin-Tyramide solution for 2 min at RT. To quench the reaction, 500 μ sodium ascorbate 3 × 5 min was added to the cells followed by 3 × 10 min washes with PBS-T. cells were lysed in 200 μ L PBS-T + 2% SDS + 2% deoxycholate + 1X complete protease inhibitor. The cells were collected by scraping, sonicated for 10 seconds, and boiled at 95°C for 60 minutes. Samples were cleared by centrifugation at maximum speed for 10 min. The supernatants were diluted in 1 mL PBS-T and 50 µL saved as input. 20 µL of pre-washed Streptavidin Dynabeads (11205D, Thermo Fisher Scientific) was added to each sample and incubated for 48 hours, rotating at 4°C. Beads were washed in 1) 15 mL PBS-T, 2) 15 mL PBS-T + 1M NaCl, 3) 15 mL PBS, and 4) 15 mL PBS+0.5% Triton X-100. Beads were re-suspended in PBS and proteins eluted by adding 1 volume of 2X lysis buffer (4% SDS, 200 mM DTT, 125 mM Tris HCl pH 6.8) and boiling at 95°C for 5 min. The eluate was separated from beads using Costar Spin-X filter units.

### Sample preparation for LC-MS/MS

Eluted proteins from light and heavy samples prepared with method 1 were mixed at a 1:1 ratio and added onto 10-kDa cutoff-filter (OD010C33, PALL) followed by addition of 1 µL 1mM DTT. Eluates prepared by method 2 were directly added onto the cut-off filters. Proteins were digested with trypsin overnight at 37°C using filter-aided sample preparation (FASP) [47]. Peptide concentration after elution was measured at 280 nm using NanoDrop (Thermo Fisher Scientific) and peptides cleaned with StageTip C18 columns [48] prior to mass-spectrometry (MS) analysis.

Eluates from proximity labeling experiments were reduced at 37°C for 60 min with DL-dithiothreitol (DTT) at 100 mM final concentration and further processed using the modified filter-aided sample preparation (FASP) method. In short, the samples were diluted 1:4 v/v by 8M urea solution, transferred onto Microcon-30kDa centrifugal units (Merck Millipore, Carrigtwohill, Ireland), and washed with 8 M urea and with Digestion buffer (0.5% sodium deoxycholate (SDC) in 50 mM TEAB. Free cysteine residues were modified using 10 mM methyl methanethiosulfonate (MMTS) solution in Digestion buffer for 20 min at room temperature and the filters were washed twice with 100 µl of Digestion buffer. Proteins were digested overnight at 37°C by adding 0.3 µg of Pierce trypsin protease (MS grade, Thermo Fisher Scientific), followed by a second incubation with 0.3 µg trypsin for three hours.

Peptides were collected by centrifugation and labelled using Tandem Mass Tag 10plex reagent (90061, Thermo Fischer Scientific) according to the manufacturer’s instructions. The labelled samples were combined into one pool, concentrated using vacuum centrifugation, and SDC was removed by acidification with 10% TFA and subsequent centrifugation. The digested peptides were cleaned-up using the HiPPR detergent removal resin kit (PN 88305, Thermo Fisher Scientific, Waltham, MA, USA) according to the manufacturer’s instructions. The sample was subsequently separated into five fractions on Pierce High pH Reversed-Phase spin column kit (Thermo Fisher Scientific) using stepwise elution with 0.1% aqueous trimethylamine solution containing 10% to 50.0% of acetonitrile. The fractions were dried and reconstituted in 15µl of 3% acetonitrile, 0.2% formic acid for LC-MS/MS analysis.

### Liquid Chromatography-MS/MS

Nano LC-MS/MS for SILAC samples was performed on a Q-Exactive HF mass-spectrometer (Thermo Fischer Scientific), connected with an EASY-nLC 1000 system (Thermo Fischer Scientific) through a nanoelectrospray ion source. Peptides were loaded on a reverse-phase column (150 mm^3^ 0.075 mm inner diameter, New Objective, New Objective, Woburn, MA) packed in-house with Reprosil-Pur C18-AQ 3 mm particles (Dr. Maisch, Ammerbuch, Germany). Peptides were separated with a 50-minute gradient: from 5 to 30% B in 35 min, 30 to 45% B in 5 min, 45 to 100% B in 1 min, followed 9 min wash with 100% of B (A: 0.1% formic acid, B: 0.1% formic acid/80% acetonitrile) using a flow rate of 250 nl/min. Q-Exactive HF was operated at 250°C capillary temperature and 2.0 kV spray voltage. Full mass spectra were acquired in the Orbitrap mass analyzer over a mass range from m/z 350 to 1600 with resolution of 60,000 (m/z 200) after accumulation of ions to a 3 × e^6^ target value based on predictive AGC from the previous full scan. Fifteen most intense peaks with a charge state ≥ 2 were fragmented in the HCD collision cell with normalized collision energy of 27%, and tandem mass spectrum was acquired in the Orbitrap mass analyzer with resolution of 15,000 after accumulation of ions to a 1 × e^5^ target value. Dynamic exclusion was set to 20 s. The maximum allowed ion accumulation times were 20 ms for full MS scans and 50 ms for tandem mass spectrum.

TMT labelled fractions were analyzed on an Orbitrap Fusion Lumos Tribrid mass spectrometer interfaced with an Easy-nLC 1200 liquid chromatography system (both Thermo Fisher Scientific). Peptides were trapped on an Acclaim Pepmap 100 C18 trap column (100 μ m, Thermo Fischer Scientific) and separated on an analytical column (75 µ m × 35 cm, packed in-house with Reprosil-Pur C18, particle size 3 µm, Dr. Maisch, Ammerbuch, Germany) using a linear gradient from 5% to 33% B over 77 min followed by an increase to 100% B for 3 min, and 100% B for 10 min at a flow of 300 nL/min. Solvent A was 0.2% formic acid in water and solvent B was 80% acetonitrile, 0.2% formic acid. MS scans were performed at 120,000 resolution in the m/z range 375-1375. The most abundant doubly or multiply charged precursors from the MS1 scans were isolated using the quadrupole with 0.7 m/z isolation window with a “top speed” duty cycle of 3 s and dynamic exclusion within 10 ppm for 45 seconds. The isolated precursors were fragmented by collision induced dissociation (CID) at 35% collision energy with the maximum injection time of 50 ms and detected in the ion trap, followed by multinotch (simultaneous) isolation of the top 10 MS2 fragment ions within the m/z range 400-1400, fragmentation (MS3) by higher-energy collision dissociation (HCD) at 65% collision energy and detection in the Orbitrap at 50,000 resolution, m/z range 100-500 and maximum injection time 105 ms.

### MS data analysis

MS raw files from SILAC experiments were processed with MaxQuant software version 1.5.7.4 [49], peak lists were identified by searching against the human UniProt protein database (downloaded 2019.04.16) supplemented with an in-house database containing all the human mucin sequences (http://www.medkem.gu.se/mucinbiology/databases/). Searches were performed using trypsin as an enzyme, maximum 2 missed cleavages, precursor tolerance of 20 ppm in the first search used for recalibration, followed by 7 ppm for the main search and 0.5Da for fragment ions. Carbamidomethylation of cysteine was set as a fixed modification. Methionine oxidation, protein N-terminal acetylation and 3-(carbamidomethyl-thio)propanoyl (DSP-crosslinker) were set as variable modifications. Arg6 and Lys6 were chosen as label modifications. The required false discovery rate (FDR) was set to 1% both for peptide and protein levels and the minimum required peptide length was set to seven amino acids.

SILAC data was analyzed with Perseus (version 1.5.5.0). First, proteins identified in the decoy database were removed together with proteins only identified by site and common contaminants. Heavy and light intensities were log_2_ transformed and filtered based on valid values in at least one group (heavy or light). Missing values were imputed based on the normal distribution of measured values using default values (width=0.3 and downshift=1.8). Significantly enriched proteins were determined with a two-sided t-test and permutation-FDR =0.05, S0=0.1 and 250 randomizations. These proteins were also manually validated as previously described [50].

Identification and relative quantification of TMT samples from proximity labelling was performed using Proteome Discoverer version 2.4 (Thermo Fisher Scientific). The database search was performed using the Mascot search engine v. 2.5.1 (Matrix Science, London, UK) against the Swiss-Prot Homo sapiens database. Trypsin was used as a cleavage rule with no missed cleavages allowed; methylthiolation on cysteine residues, TMT at peptide N-termini and on lysine side chains were set as static modifications, and oxidation on methionine was set as a dynamic modification. Precursor mass tolerance was set at 5 ppm and fragment ion tolerance at 0.6 Da. Percolator was used for the peptide-spectrum match (PSM) validation with the strict false discovery rate (FDR) threshold of 1%. Quantification was performed in Proteome Discoverer 2.4. The TMT reporter ions were identified with 3 mmu mass tolerance in the MS3 HCD spectra and the TMT reporter S/N values for each sample were normalized within Proteome Discoverer 2.4 on the total peptide amount. Only the unique identified peptides were considered for the protein quantification.

### Immunofluorescence and image analysis

Staining of Caco-2 cells grown on chamber slides (154534PK, Thermo Fisher Scientific) were performed as described previously [24]. In brief, cells were washed 3 times in warm PBS followed by 10 min fixation in 4% paraformaldehyde (PFA) in PBS at RT. Excess PFA was washed away with 3 PBS washes and cells permeabilized by 0.1% Triton X-100 in PBS for 7 min. After permeabilization, cells were washed three times with PBS and blocked overnight in 5% BSA in PBS at 4°C. Cells were incubated with primary antibodies diluted in 5% BSA in PBS for 2 hours at 24°C, then washed 3 times with PBS and incubated with secondary antibodies for 1 hour at RT. After three washes with PBS, cell nuclei were stained with Hoechst for 7 min at RT. Chamber slides were washed 3 times with PBS and mounted with Prolong anti-fade (Invitrogen).

Harvested ileum was fixed in Carnoy’s fixative (60% absolute methanol, 30% chloroform and 10% glacial acetic acid) or 4% paraformaldehyde (PFA) solution. Samples fixed in Carnoy’s fixative were embedded in paraffin. Paraffin-embedded sections were deparaffinized in xylene substitute (2 3 10 min, 60°C) and rehydrated in 100% ethanol (10 min), 70% (v/v) ethanol (5 min), 50% (v/v) ethanol (5 min), and 30% (v/v) ethanol (5 min). Sections were placed in antigen retrieval buffer (0.01M citric acid, pH 6.0) at 100 degrees for 10 min and then brought to RT (2 hours) and transferred to PBS. Tissues were enclosed with a PAP pen followed and blocked with 5% fetal calve serum (FCS) in PBS for 20 min at RT. Primary antibodies were diluted in 5% FCS in PBS and incubated overnight at 4°C. After 3x5 min washes in PBS, sections were incubated with secondary antibodies diluted in 5% FCS in PBS. DNA was stained with Hoechst for 5 min at RT. Coverslips were mounted using Prolong Gold antifade (P36980, ThermoFisher Scientific) and polymerized overnight at RT in the dark. Images and Z-stacks were acquired on a Zeiss LSM 700 (Plan-Apochromat 40x/1.3 Oil DIC M27 lens and 1.58µs pixel dwell) and a Zeiss LSM900 (equipped with an Airyscan2 detector and plan-Apochromat 63x/1.4 Oil DIC M27 lens). All image analysis and processing were performed in ImageJ software v.1.53.t (National Institutes of Health, Bethesda, MD). Confocal images are shown as maximum projections except from YZ and XZ sections. Line intensity profiles were generated from z-axis profiles for each individual channel, averaged to 15 µm distance from the basolateral to apical membrane and normalized to values between 0-100.

### Transmission electron microscopy

MUC17(7TR) Caco-2 cells grown on Transwell filters (CLS3496, Merck) for 7, 14 and 21 DPC were fixed in primary fixative (1.33% glutaraldehyde in water, 0.1 M cacodylate buffer, 0.05% Ruthenium Red in water) for 1 hour at RT. Fixed cells were pre-washed in 0.05 M cacodylate buffer and then washed extensively in 2x10 min 0.05 M cacodylate buffer, 20 min 0.05 M cacodylate buffer + 0.02 M glycine followed by 2x10 min 0.05 M cacodylate buffer. Secondary fixative (1.33% osmium tetroxide in water, 0.1 M cacodylate buffer, 0.05% Ruthenium Red in water) was added to the cells for 1 hour and incubated at 4 °C in the dark on an orbital shaker. Secondary fixative was removed by a few quick washes in water followed by 6x5 min washes in water. Cells were incubated with tertiary fixative (1% filtered aqueous uranyl acetate) for 30 min at room temperature in the dark followed by a few washes with water. Excised membranes were stained in lead aspartate (0.02 M lead nitrate and 0.03M aspartic acid pH 5.5) for 20 min at RT. Cells were washed three times in water, incubated overnight in water followed by another three washes in water. Dehydration of cells was done in a series of ethanol solutions at RT (5 min 30% EtOH, 5 min 50% EtOH, 5 min 70% EtOH, 5 min 85% EtOH, 5 min 95% EtOH, 5x5 min 100% EtOH). The cells were embedded in Hard-Plus Epoxy 812-resin (14115, Electron Microscopy Sciences, US) at RT as follows. 25% resin in acetone for 1 hour, 50% resin 2 hours, 75% resin 1 hour, 100% resin 3x30 min and 100% resin overnight on an orbital shaker. After another incubation of 2 × 1 hour 100% resin, samples were incubated with 100% resin + accelerator (240 µL/10 mL) for 1 hour and another 100% + accelerator for several hours. Infiltrated samples were embedded in resin-silicon free molds with resin + accelerator and let to polymerize for 48 hours at 60°C. Transversal ultrathin sectioning (70-90 nm) of cells from the apical to basolateral membrane were performed on Leica UC6 Ultracut, collected on copper 100 hexagonal mesh support grids and post stained with Reynold’s solution [51] at RT for 5 min in a sealed chamber with NaOH-pellets. Sections were imaged on TEM FEI Talos (ThermoFisher, US) equipped with 4kx4k Ceta CMOS camera operating at 120 kV with LaB6 filament.

### Fluorescence Recovery After Photobleaching (FRAP)

StcE E447D was labeled with CF 555 Succinimidyl Ester (92214, Biotium) according to manufactureŕs protocol. 500 µL serum-free medium containing 10µg/mL of CF555-StcE E447D (RT) was added to WT and knockout Caco-2 cells differentiated on 30 mm plates for 30 min. After two washes with 3 mL PBS (RT), images were acquired using LSM700 with a plan-apochromat × 20/1.0 DIC M27 75 mm water objective (Zeiss) at 1.5 digital zoom and ZEN 2010 software. Photobleaching at 100% laser power for a duration of 25 µsec was performed after 5 initial scans (pixel dwell per scan) using the 555 nm laser. Four 20 µm × 20 µm square regions of interest (ROIs) where selected according to the following scheme: ROI1: analyze in cell 1 (bleach control) and ROI2: bleach and analyze in cell 2. Recovery images were acquired every 2 seconds during 12 min. Raw data were analyzed by the formula,

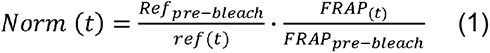

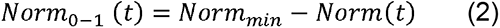

Where Ref_pre-bleach_ is the mean intensity of ROI1 before bleaching, FRAP_pre-bleach_ is the mean intensity of the ROI2 pre bleaching, ref(t) is the intensity of ROI1 at time point t, FRAP(t) in the intensity of ROI2 at time point t. Norm(t) represent fluorescence in ROI2 at time (t) corrected for bleaching during analysis recovery. Norm_0-1_(t) sets the mean intensity of ROI2 before bleaching to 1 and after bleaching to 0. Recovery halftimes and mobile fractions were extracted by fitting the data to a non-linear curve based on a one-phase association.

### Quantification and statistical analysis

Data analysis was performed using GraphPad Prism (version 9.5) and Perseus (version 1.5.5.0). Graphs were prepared using either Perseus (version 1.5.5.0) or GraphPad Prism (version 9.5). Venn Diagrams were created using: https://bioinformatics.psb.ugent.be/webtools/Venn/. One- or two-way ANOVA followed by Tukeýs or Sidak’s multiple comparisons test or Kruskal Wallis and Dunn’s multiple comparisons test was done for comparisons of multiple groups. Unpaired t-test with Welch’s correction, assuming non-equal SDs was used for comparison of two groups. *p< 0.05, **p<0.01, ***p<0.001, ****p<0.0001.

## Author contributions

Conceptualization, S.J. and T.P.; methodology, S.J. and T.P.; investigation, S.J., G.H., I.K., and T.P.; writing – original draft, S.J. and T.P.; writing – review & editing, S.J., I.K. J.R.G., and T.P.; funding acquisition, S.J., I.K., J.R.G., and T.P.; resources, T.P.; supervision, T.P.

## Acknowledgements

We acknowledge Drs. Carina Sihlbom, Egor Vorontsov, and Evelin Berger at the Proteomics Core Facility of Sahlgrenska Academy (University of Gothenburg) for performing TMT proteomic analysis. We thank colleagues at the Centre for Cellular Imaging Core Facility of Sahlgrenska Academy (University of Gothenburg) for sample preparation and image acquisition for transmission electron microscopy. This study was supported by the Swedish Society for Medical Research (grant S17-0005) to T.P., National Institutes of Health (NIH) (grants 5U01AI095542-08-WU-19-95 and 5U01AI095542-09-WU-20-77) to T.P., NIH (grants RC2 DK118640 and R01 DK48370) to J.R.G., NIH (grant R01 DK128190) to I.K., Wenner-Gren Foundations (grant FT2017-0002) to T.P., Jeansson Foundations (grant JS2017-0003) to TP, Åke Wiberg Foundation (grant M17-0062) to T.P., Stiftelsen Clas Groschinskys Minnesfond (grant M2254) to T.P., and Sahlgrenska Academy doctoral project grant (grant U2018/162) to S.J. and T.P..

