## Supplementary material for "The MYO1B and MYO5B motor proteins and the SNX27 sorting nexin regulate membrane mucin MUC17 trafficking in enterocytes"

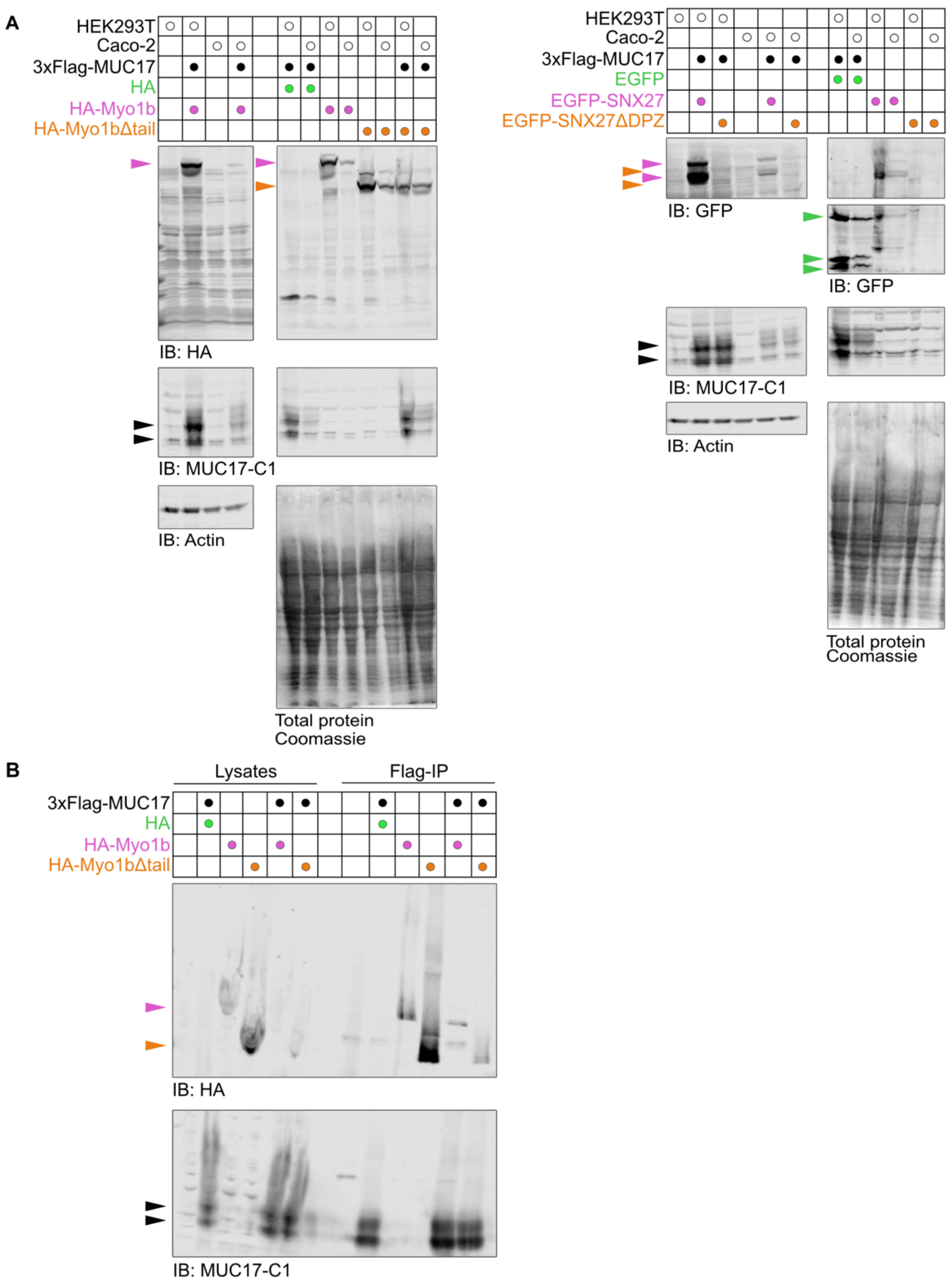


**Figure S1.**

A) Validation of recombinant protein expression in stable HEK293T and Caco-2 clones. Immunoblots of HEK293T and Caco-2 cells expressing combinations of 3xFlag-MUC17(7TR), HA, HA-Myo1b and HA-Myo1bΔtail (left panels) or 3xFlag-MUC17(7TR), EGFP, EGFP-SNX27 and EGFP-SNX27ΔPDZ (right panels).

B) Representative immunoblot of co-immunoprecipitations in HEK293 cells expressing 3xFlag-MUC17(7TR) and either HA-Myo1b or HA-Myo1bΔtail. Lysates represent 2% of the total cell lysate whereas 40% of the eluates was loaded on the gel.

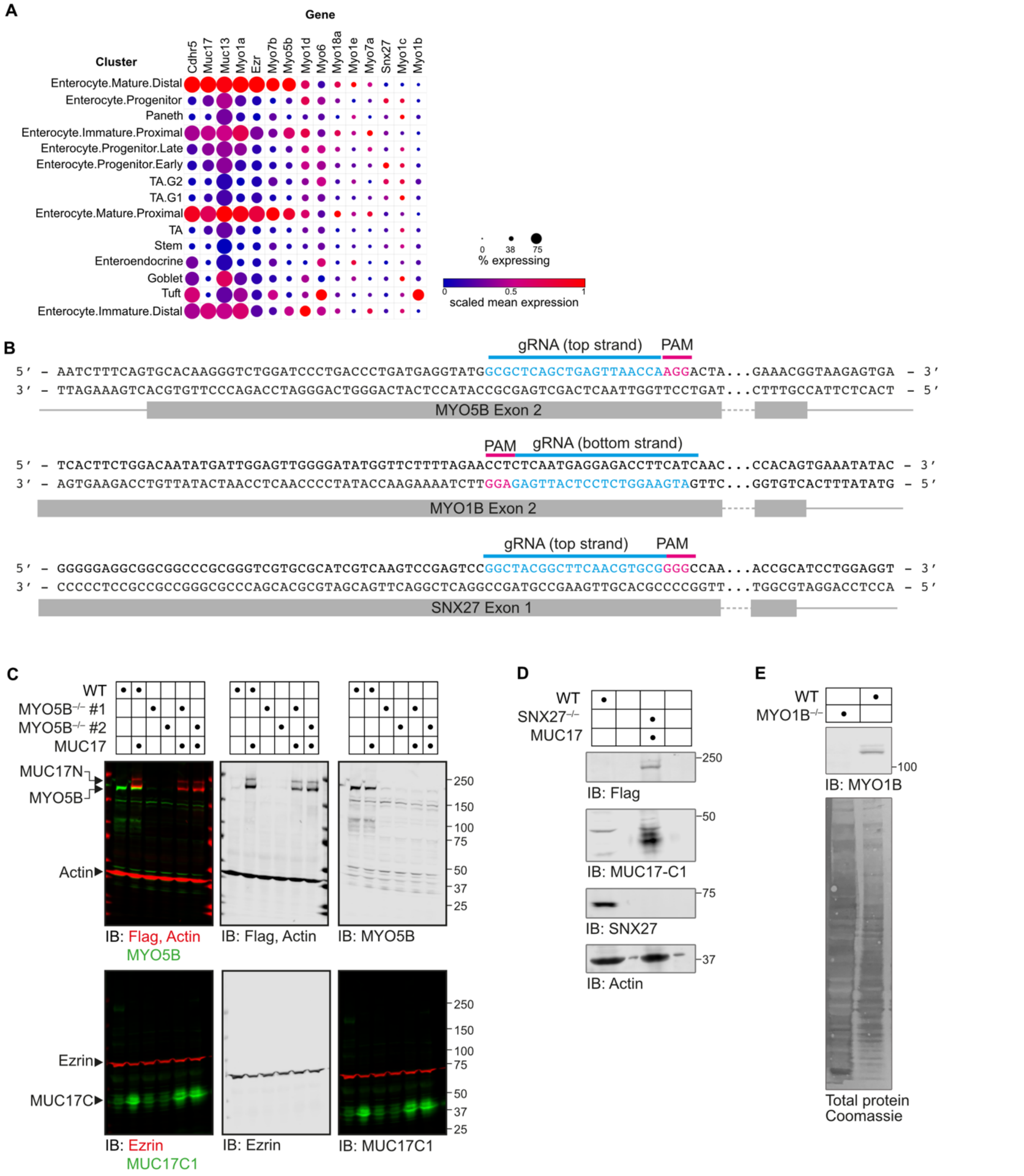


**Figure S2.**

A) Single-cell transcriptomics from murine small intestine [1] showing non-muscle myosins and core microvillar components identified in [2]. Graphical visualization was performed using the Broad Institute Single Cell Portal.

B) Placement of Crispr/Cas9 guide RNAs for deletion of human *MYO5B*, *MYO1B* and *SNX27* genes.

C) Validation of Crispr/Cas9-mediated deletion of *MYO5B*, *SNX27* (D), and *MYO1B* (E) in Caco-2 cells.

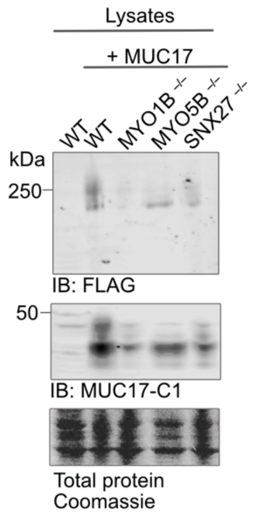


**Figure S3.**
Immunoblot of total MUC17(7TR) and total protein for WT and KO cells used for quantification in figure 7A.

**A  Optimized sequence of cDNA insert encoding 3xFlag-MUC17(7TR)**

```
ATGCCTAGACCTGGGACTATGGCATTGTGCCTCCTTACACTGGTGCTGTCTCTCCTCCCTCCTCAGGCTGCCGCTGAG
CAGGATCTCAGCGTGAACAGAGCAGTGTGGGATGGAGGAGGCTGTATTTCTCAAGGGGATGTGCTGAATCGGCAGTGT
CAGCAACTGAGCCAGCATGTACGCACAGATTACAAAGACCACGACGGCGACTACAAGGATCACGACATCGACTACAAG
GACGACGACGACAAAGGCGAGAATCTGTATTTCCAGTCTGGCGGGGGGTCAGCTGCCAACACAGCAACAGGGACAACC
TCCACAAATGTGGTGGAACCCCGAATGTATCTGAGCTGTAGCACAAACCCTGAAATGACCTCCATTGAGTCAAGCGTC
ACATCAGACACACCGGGTGTTTCCAGTACGCGCATGACTCCAACCGAGTCACGTACCACGAGCGAGTCTACTTCAGAT
TCTACAACCCTCTTCCCCAGTAGTACCGAGTCCTCTAGCCCTCCTATTGCTGACGGAACCAGTATGCCAACGTCTACG
TATAGTGAAGGTTCTACTCCCTTAACAAACATGTCATTTAGTACAACTCCAGTCGTTAGTAGCGAGGCGAGTACATTA
TCTACAACCCCCGTGGACACATCGACGCCTGTGACCACGAGCACGGAAGCATCCCTAAGTCCTACTACTGCAGAAGGG
ACCAGTATTCCTACATCGTCACCATCTGAGGGACAACGCCACTGGCTTCTATGCCCGTCAGCACTACGCCAGTTGTA
TCGAGCGAAGTTAACACATTGTCAACGACGCCCGTGGATTCCAACACACTGGTGACCACCTCCACTGAAGCAAGCTCC
TCGCCGACCATTGCAGAAGGAACTTCACTGCCGACAAGTACAACTTCGGAGGGATCAACACCGTTGTCTATCATGCCC
CTCTCCACCACACCGGTGGCCTCTTCTGAGGCCAGCACACTCTCAACTACTCCGGTGGACACCTCTACCCCCGTCACC
ACCTCTTCACCAACTAACTCTTCTCCTACTACCGCTGAGGTGACCTCTATGCCAACTAGCACCGCGGGCGAGGGCAGT
ACCCCACTCACCAATATGCCTGTATCCACCACTCCCGTTGCTTCCTCTGAGGCTAGTACACTCAGCACGACTCCTGTT
GACAGCAATACCTTCGTGACTAGCTCATCTCAGGCCTCATCCTCACCTGCCACACTGCAGGTCACAACTATGAGGATG
TCCACCCCTAGTGAGGGTAGTTCCTCCCTTACCACTATGCTTCTGTCAAGTACCTACGTGACCAGCAGCGAGGCCTCC
ACACCCAGTACCCCTAGCGTGGATAGAAGCACCCCAGTCACAACTAGCACTCAGAGCAATTCCACTCCAACCCCACCA
GAAGTGATCACCCTGCCCATGTCCACCCCCAGCGAGGTAAGCACTCCCCTGACAATCATGCCCGTGTCCACCACCTCC
GTCACAATCTCCGAAGCCGGCACAGCCTCCACCCTGCCCGTCGATACCTCCACACCTGTGATAACATCCACCCAAGTG
AGTTCATCTCCTGTGACTCCTGAAGGTACCACCATGCCAATCTGGACGCCTAGTGAAGGAAGCACTCCATTAACAACT
ATGCCTGTCAGCACCACACGTGTGACCAGCTCTGAGGGTAGCACCCTTTCAACACCTTCTGTTGTCACCAGCACACCT
GTGACCACTTCTACTGAAGCCATTTCATCTTCTGCAACTCTTGACAGCACCACCATGTCTGTGTCAATGCCCATGGAA
ATAAGCACCCTTGGGACCACTATTCTTGTCAGTACCACACCTGTTACGAGGTTTCCTGAGAGTAGCACCCCTTCCATA
CCATCTGTTTACACCAGCATGTCTATGACCACTGCCTCTGAAGGCAGTTCATCTCCTACAACTCTTGAAGGCACCACC
ACCATGCCTATGTCAACTACGAGTGAAAGAAGCACTTTATTGACAACTGTCCTCATCAGCCCTATATCTGTGATGAGT
CCTTCTGAGGCCAGCACACTTTCAACACCTCCTGGTGATACCAGCACACCTTTGCTCACCTCTACCAAAGCCGGTTCA
TTCTCCATACCTGCTGAAGTCACTACCATACGTATTTCAATTACCAGTGAAAGAAGCACTCCATTAACAACTCTCCTT
GTCAGCACCACACTTCCAACTAGCTTTCCTGGGGCCAGCATAGCTTCGACACCTCCTCTTGACACAAGCACAACTTTT
ACCCCTTCTACTGACACTGCCTCAACTCCCACAATTCCTGTAGCCACCACCATATCTGTATCAGTGATCACAGAAGGA
AGCACACCTGGGACAACCATTTTTATTCCCAGCACTCCTGTCACCAGTTCTACTGCTGATGTCTTTCCTGCAACAACT
GGTGCTGTATCTACCCCTGTGATAACTTCCACTGAACTAAACACACCATCAACCTCCAGTAGTAGTACCACCACATCT
TTTTCAACTACTAAGGAATTTACAACACCCGCAATGACTACTGCAGCTCCCCTCACATATGTGACCATGTCTACTGCC
CCCAGCACACCCAGAACAACCAGCAGAGGCTGCACTACTTCTGCATCAACGCTTTCTGCAACCAGTACACCTCACACC
TCTACTTCTGTCACCACCCGTCCTGTGACCCCTTCATCAGAATCCAGCAGGCCGTCAACAATTACTTCTCACACCATC
CCACCTACATTTCCTCCTGCTCACTCCAGTACACCTCCAACAACCTCTGCCTCCTCCACGACTGTGAACCCTGAGGCT
GTCACCACCATGACCACCAGGACAAAACCCAGCACACGGACCACTTCCTTCCCCACGGTGACCACCACCGCTGTCCCC
ACGAATACTACAATTAAGAGCAACCCCACCTCAACTCCTACTGTGCCAAGAACCACAACATGCTTTGGAGATGGGTGC
CAGAATACGGCCTCTCGCTGCAAGAATGGAGGCACCTGGGATGGGCTCAAGTGCCAGTGTCCCAACCTCTATTATGGG
GAGTTGTGTGAGGAGGTGGTCAGCAGCATTGACATAGGGCCACCGGAGACTATCTCTGCCCAAATGGAACTGACTGTG
ACAGTGACCAGTGTGAAGTTCACCGAAGAGCTAAAAAACCACTCTTCCCAGGAATTCCAGGAGTTCAAACAGACATTC
ACGGAACAGATGAATATTGTGTATTCCGGGATCCCTGAGTATGTCGGGGTGAACATCACAAAGCTACGTCTTGGCAGT
GTGGTGGTGGAGCATGACGTCCTCCTAAGAACCAAGTACACACCAGAATACAAGACAGTATTGGACAATGCCACCGAA
GTAGTGAAAGAGAAAATCACAAAAGTGACCACACAGCAAATAATGATTAATGATATTTGCTCAGACATGATGTGTTTC
AACACCACTGGCACCCAAGTGCAAAACATTACGGTGACCCAGTACGACCCTGAAGAGGACTGCCGGAAGATGGCCAAG
GAATATGGAGACTACTTCGTAGTGGAGTACCGGGACCAGAAGCCATACTGCATCAGCCCCTGTGAGCCTGGCTTCAGT
GTCTCCAAGAACTGTAACCTCGGCAAGTGCCAGATGTCTCTAAGTGGACCTCAGTGCCTCTGCGTGACCACGGAAACT
CACTGGTACAGTGGGGAGACCTGTAACCAGGGCACCCAGAAGAGTCTGGTGTACGGCCTCGTGGGGGCAGGGGTCGTG
CTGATGCTGATCATCCTGGTAGCTCTCCTGATGCTCGTTTTCCGCTCCAAGAGAGAGGTGAAACGGCAAAAGTACAGA
TTGTCTCAGTTATACAAGTGGCAAGAAGAGGACAGTGGACCAGCTCCTGGGACCTTCCAAAACATTGGCTTTGACATC
TGCCAAGATGATGATTCCATCCACCTGGAGTCCATCTATAGTAATTTCCAGCCCTCCTTGAGACACATAGACCCTGAA
ACAAAGATCCGAATTCAGAGGCCTCAGGTAATGACGACATCATTTTAA
```

**Color codes:**
Startcodon – 3xFlag – FlexLink – PTS1 – PTS 55-60 – SEA - TM

B

| Forward and reverse primers used to generate Endo-SS-3xFlag-MUC17(7TR) | |
|---|---|
| Kozak- endo-SS -3xFlag-PTS1 | 5´- CAGGGCCAGATATCGCCGCCACCATGCCTAGACCTGGGACTATG - 3´<br>5´- AATAGGAGGGCTAGAGGAGTC-CTCGGTACTACTGGG - 3´ |
| TR 1-6 | 5´- CCCAGTAGTACCGAGGACTCCTCTAGCCCTCCTATT - 3´<br>5´- ATAACATCCACCCAAGTGAGTTCATCTCCTGTGACT - 3´ |
| SEA-TM-CT | 5´- ATAACATCCAC-CCAAGTGAGTTCATCTCCTGTGACT - 3´<br>5´- TGAGCTTTTGCTCTGGTTAAAATGATGTCGTCATTACCTGA - 3´ |

**Figure S4.**
A) DNA sequence for generating 3xFlag-MUC17(7TR).

B) Forward and reverse primer combinations for generation of 3xFLAG-MUC17(7TR) with overhangs for the pXL-CAG-Zeocin-3xF2A plasmid.
